# Timely lagging strand maturation relies on Ubp10-mediated PCNA dissociation from replicating chromatin

**DOI:** 10.1101/2024.01.05.574312

**Authors:** Javier Zamarreño, Sofía Muñoz, Esmeralda Alonso, Macarena Alcalá, Rodrigo Bermejo, María P. Sacristán, Avelino Bueno

**Author notes:** JZ, SM and EA, contributed equally to this work.

## Abstract

Synthesis and maturation of Okazaki Fragments is an incessant and highly efficient metabolic process completing the synthesis of the lagging strands at replication forks during S phase. Accurate Okazaki fragment maturation (OFM) is crucial to maintain genome integrity and, therefore, cell survival in all living organisms. In eukaryotes, OFM involves the consecutive action of DNA polymerase Pol ∂, 5’ Flap endonuclease Fen1 and DNA ligase I, and constitutes the best example of a sequential process coordinated by the sliding clamp PCNA. For OFM to occur efficiently, cooperation of these enzymes with PCNA must be highly regulated. Here, we present evidence of a role for the PCNA-deubiquitylase Ubp10 in the maturation of Okazaki fragments in the budding yeast *Saccharomyces cerevisiae*. We show that Ubp10 associates with lagging-strand DNA synthesis machineries on replicating chromatin to ensure timely ligation of Okazaki fragments by promoting an Elg1^ATAD5^-independent PCNA unloading mechanism.

This document was written without the use of AI.

## INTRODUCTION

The *POL30* gene of *Saccharomyces cerevisiae* encodes the sliding clamp PCNA (Proliferating Cell Nuclear Antigen), a conserved ring-shaped protein with essential roles in DNA metabolism as a crucial component of replication and repair machineries. PCNA forms an homotrimer that encircles DNA, where it interacts with a staggering number of proteins involved in every step required for DNA replication or repair. Thus, whereas it is devoid of enzymatic activity itself, PCNA exerts its function by recruiting and, in many instances, also activating numerous interactors^1,2^.

DNA polymerases require free 3’-OH groups to initiate DNA synthesis, therefore, they can only synthesize DNA in a 5’-3’ direction. As a result of this and the antiparallel nature of double stranded DNA, replication of one of the strands is discontinuous through the generation of Okazaki Fragments (OF). This strand is called the lagging strand (in contrast to the leading one). PCNA plays a crucial role in the synthesis and maturation of these OFs. Thus, PCNA is loaded on dsDNA at primer-template junctions to recruit Pol δ and enhance its processivity. When the Pol δ-PCNA complex collides with the 5’-end of the preceding OF, it displaces a short flap that is cleaved off by the structure-specific flap endonuclease-1 Fen1, Rad27 in *S. cerevisiae*, upon binding to PCNA. This process generates a nick in the nascent DNA that is sealed by DNA ligase I, Cdc9 in *S. cerevisiae*, which is also recruited and catalytically activated through its interaction with PCNA^3^. All three subunits of DNA Pol δ, Fen1/Rad27, and DNA ligase I/Cdc9 harbor PIP (PCNA-interacting peptide)-boxes, through which they interact in a coordinated manner with the Inter-Domain Connecting Loop (IDCL) of PCNA, a major interaction site in the sliding clamp^4–6^. Although the mechanisms and factors involved in lagging strand maturation have been extensively studied key molecular details of OF maturation remain poorly understood.

To ensure the successful completion of DNA replication, particularly of the processive synthesis of the lagging strands, continuous recycling of chromatin-bound PCNA is required. The sliding clamp is loaded when required and actively unloaded when no longer needed in order to suppress illegitimate enzymatic reactions^2,7,8^. RFC (Replication factor C) and RFC-like complexes (RLCs) mediate the loading and unloading of PCNA (reviewed in^9^). There is a general consensus regarding Rfc1-RFC complex role as the main loader of PCNA on replicating chromatin^10,11^. One additional complex, Ctf18-RLC, also acts as a PCNA loader, although Ctf18 cannot substitute Rfc1. Moreover, Rfc1 and Ctf18 show a strand preference during replication, with a lagging strand bias for Rfc1 and a slight leading strand preference for Ctf18^12–14^. Both Rfc1 and Ctf18 also exhibit some ability to unload PCNA *in vitro*. Human Rfc1-RFC, but not yeast Rfc1, is able to unload PCNA *in vitro* in an ATP-dependent manner^15,16^ and yeast Ctf18-RLC promotes an *in vitro* PCNA unloading mechanism, that also requires ATP hydrolysis, in the presence of ssDNA coated with RPA^17^. Despite these *in vitro* results, PCNA unloading activities for Rfc1- or Ctf18-complexes *in vivo* have not been reported.

Elg1-RLC complex (ATAD5-RLC in mammals) is considered as the major PCNA unloader when the role of PCNA in DNA replication is completed^8,18–22^. However, given that *ELG1* is a non-essential gene for cell division^23–25^ and that PCNA accumulated in Elg1-depleted cells ends up being removed from chromatin before M phase^18,20^, it is likely that additional PCNA unloaders are required during chromosome replication, at least in the absence of Elg1. Therefore, it remains unclear whether Elg1-RLC is the only in vivo PCNA unloader^22,26^.

PCNA functions are regulated by different post-translational modifications such as SUMOylation, ubiquitylation, phosphorylation or acetylation, which confer PCNA the necessary plasticity to interact with its different binding partners^1,27,28^. In the face of DNA lesions, PCNA is ubiquitylated to mediate damage-tolerance mechanisms that allow circumventing DNA lesions and prevent replication fork stalling^29,30^. PCNA is mono-ubiquitylated at K164 by the evolutionary conserved RAD6/RAD18 (E2/E3) ubiquitin ligase complex to switch its affinity from replicative polymerases to damage-tolerant translesion synthesis (TLS) DNA polymerases, which, although mutagenic, are capable to bypass damaged bases^27,31,32^. Furthermore, polyubiquitylation of the same residue by the Rad5/Mms2/Ubc13 PCNA-ubiquitin ligase complex leads to template switching (TS), the DNA damage tolerance (DDT) error-free pathway, to overcome the potentially lethal effects of replication fork stalling^27,30^. We and others reported that the precise regulation of these processes not only depends on writer enzymes, PCNA^K164^– Ubiquitin ligase complexes, but also on the erasers of these modifications. Thus, both TLS and TS pathways are limited by PCNA-deubiquitylation processes to minimize their deleterious cellular side effects. In mammals, deubiquitylating enzymes Usp1, Usp7, and Usp10 revert PCNA ubiquitylation caused in response to DNA damage^28,33–35^. Knockdown of USP1 induces aberrant PCNA monoubiquitylation, enhanced recruitment of error-prone TLS polymerases and increased mutagenesis levels in human cells^33,35^. In the case of the budding yeast *S. cerevisiae*, the PCNA-DUBs Ubp10 and Ubp12^36,37^ limit the extent of DDT processes during the progression of exogenously unperturbed S phase by reverting K164-ubiquitylation of the sliding clamp at replication forks^37^. Ubp10 ubiquitin-protease has also a role deubiquitylating histone H2B^K123^ working at genomics sites distinct of the SAGA-related histone H2B^K123^ DUB Ubp8 sites^38–40^. Moreover, Ubp10 activity is also involved in the regulation of RNA polymerase I stability^41^. Remarkably, despite the key roles of this PCNA-DUB, abrogation of the *UBP10* gene is viable, even though *ubp10* mutated cells have both growth and cell cycle progression defects^36,37,41,42^. Suppression of either growth or cell cycle defects can be accomplished by mutation of specific targets. Thus, multiple deletion of the TLS polymerases (*REV1*, *REV3* and *RAD30*) rescues the cell cycle delay in S phase progression caused by the abrogation of Ubp10^37^, indicating that the role of Ubp10 in supporting normal replication rates through PCNA-K164 deubiquitylation is dependent, at least in part, on the TLS pathway.

A number of active roles in the regulation of key cellular mechanisms has been described for ubiquitin-signalling writers, but not so many for erasers. Along this line of thought, it is assumed that pivotal regulatory steps rest on ubiquitin writers while redundant erasers are considered to act automatically or spontaneously after the post-translational modification is added to a given substrate. For these reasons ubiquitin proteases are in general considered to play a minor role, if any, in regulatory controls. In this context, it is assumed that PCNA ubiquitylation is counteracted by constitutive deubiquitylation mediated by redundant PCNA-DUBs^43,44^. However, recent work with yeast models concerning PCNA-DUBs role in DNA replication shows that something is amiss with this scenario both in fission and budding yeast^37,43^. Regarding this matter, Ubp10 has a remarkable slow S phase phenotype that we were very interested to understand in full.

PCNA^K164^ ubiquitylation has also been linked to the OFM process. Thus, loss of PCNA ubiquitylation seems to cause inefficient gap-filling which interferes with efficient OF ligation in fission yeast and human cells^44,45^. Moreover, PCNA^K164^ ubiquitylation suppresses replication stress resulting from Fen1/Rad27-defective flap processing during OFM^46^, and the PCNA^K164R^ mutant shows inefficient OF processing in an *in vitro* DNA replication reconstitution assay using yeast proteins^47^. Here, we present evidence of the role of Ubp10 in the regulation of OFM. Mass spectrometry analysis of the Ubp10 interactome showed that this PCNA-DUB interacts with all major components of the OF metabolism. Based on this observation and on the cell cycle delay in S phase, characteristic of cells deficient in Ubp10, we focused on deciphering the potential link between Ubp10 and OFM processes. Ablation of Ubp10 leads to accumulation of unligated OFs and a markedly increase of chromatin-bound PCNA during S phase. *POL30* mutants that conform unstable PCNA homotrimers on chromatin, particularly *pol30^R14E^* and *pol30^D150E^* alleles, counteract these *ubp10*Δ-associated replication defects, as well as cell cycle delay. In addition, abrogation of *ubp10* is strongly additive to *elg1* depletion, resulting in substantial increase of the PCNA bound to chromatin during replication. These data indicate that timely dissociation of PCNA during lagging strand synthesis requires Ubp10 in an Elg1/ATAD5 independent manner and is important for OF ligation. All this evidence reveals an important regulatory role for Ubp10 at the lagging strand synthesis.

## RESULTS

### PCNA-DUB Ubp10 physically interacts with core components of the lagging-strand synthesis machinery

We first analyzed the interactome of the H2B- and PCNA-ubiquitin protease Ubp10 to reveal partners of this DUB during DNA replication by LC-MS-MS analysis. Mass spectrometry analysis of the proteome associated with Ubp10 in unperturbed S phase cells retrieved all major components of the lagging strand synthesis machinery (such as PCNA, Polα/primase, RFC replication clamp loader, DNA polymerase ∂) and Okazaki fragment metabolism (Fen1 flap endonuclease, Cdc9 ligase, RNase H2) (Figure 1). A complementary proteomic analysis of DNA ligase Cdc9 also revealed Ubp10 as one of the DNA ligase interactors (to be published elsewhere). Relevant interactions were confirmed by direct co-immunoprecipitation (ChIP-CoIP) analyses in epitope-tagged backgrounds, as shown for Pol3, catalytic subunit of Polδ, (Supplementary Figure 1A), and Cdc9 (Supplementary Figure 1B). Ubp10-PCNA and Ubp10-Fen1 ChIP-CoIP interactions have been described previously^36,37^. This evidence suggests the existence of a functional link between Polδ, Fen1, Cdc9, Ubp10 and PCNA, and, therefore, supports the hypothesis that Ubp10 works on the lagging strand during OFM. Likely significant, our Ubp10-GFP-trap proteomic approach did not detect RLC cofactor Elg1. In contrast, all Rfc1-RFC subunits were identified as Ubp10 unperturbed S phase interactors. Of interest, FACT subunits (Spt16 and Pob3) and RNA pol I subunits (Rpa190, 34, 43, 49 and 135), comprising known functional interactors of Ubp10 Spt16 and Rpa190 ^41,48^, were identified with high scores in our proteomic analyses validating our experimental approach (Supplementary Figure 1C). (All these detected Ubp10 interactions somehow endorse our LC-MS-MS analysis).

**Figure 1.**
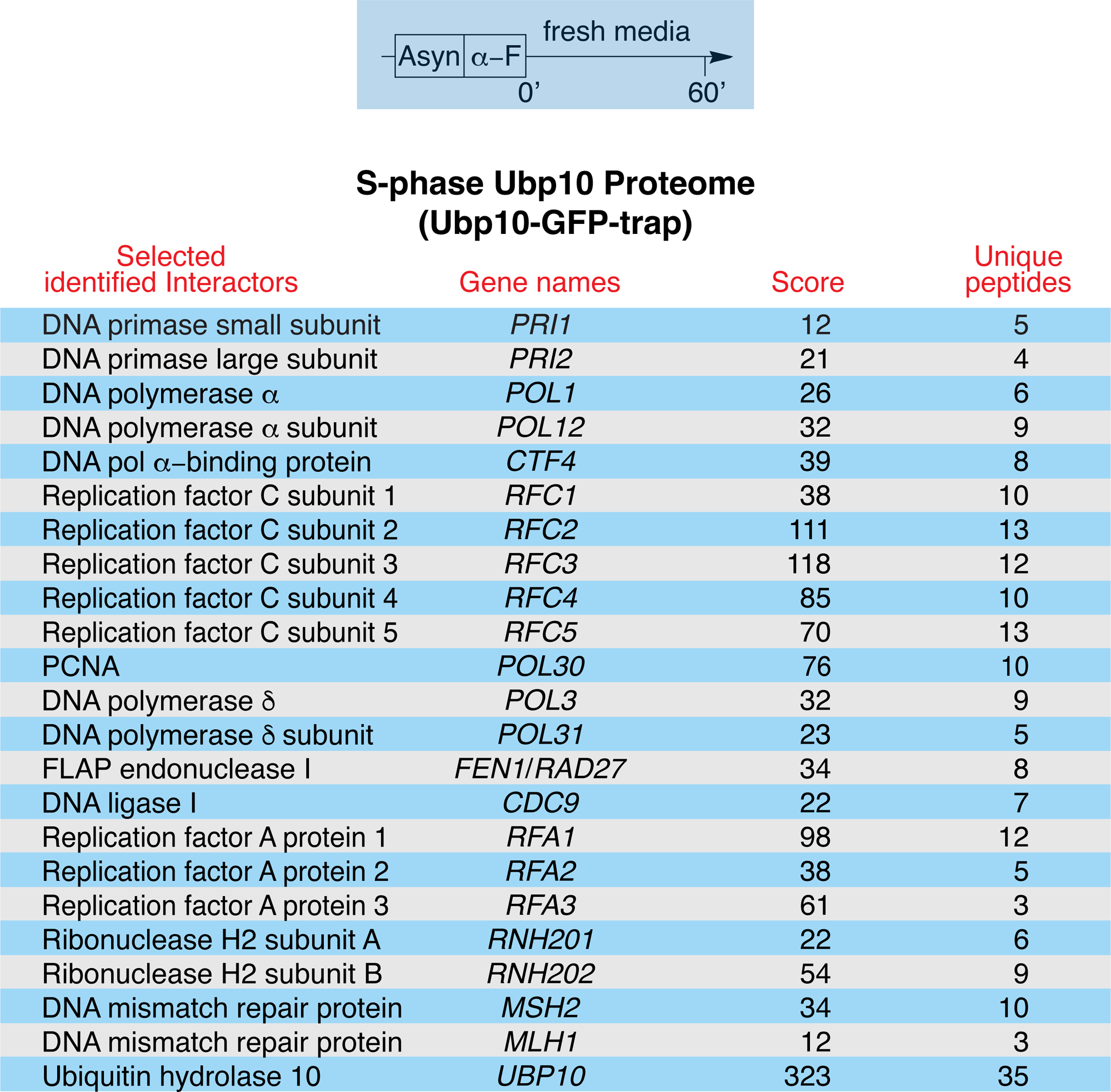
PCNA-DUB Ubp10 interacts with core components of lagging-strand synthesis machinery. PCNA-DUB Ubp10 interacts with all major lagging-strand synthesis complex components during unperturbed S phase as detected by LC-MS-MS analysis. Ubp10-GFP was immunoprecipitated from yeast cells pre-synchronized in G1 with α-factor and released 60 minutes in fresh rich media (S phase). Immunoprecipitation was carried out by using a GFP-Trap Agarose beads as described in Methods and using an untagged strain as a negative control. Full IPs were subject to LC-MS-MS analysis. The list shows Ubp10-GFP interacting proteins identified by LC-MS-MS. Potential Ubp10-interactors were identified among proteins with at least 3 unique peptides (only selected interactors are shown). Some selected proteins were chosen for confirmative CoIP analysis as shown in supplementary figure 1 for Pol3 (catalytic subunit of Pol∂) and Cdc9 (DNA ligase I) together with LC-MS-MS results obtained with other proteins of interest regarding Ubp10 biology.

### The PCNA-DUB Ubp10 acts downstream of the Fen1^Rad27^ FLAP-endonuclease during DNA replication

Our Ubp10-GFP-Trap proteomic approach shows that Ubp10 interacts physically with major components of the lagging strand synthesis machinery during S phase. Therefore, looking for a functional support of the proteomic evidence, a molecular analysis was designed to reveal a potential role of Ubp10 in the metabolism of OFs. Initially, we focused in discerning the potential interplay of *FEN1* and *UBP10* in the OFM pathway. We had recently shown that Ubp10 and the Fen1 Flap-endonuclease physically interact in early S phase, as detected by co-immunoprecipitation^37^. To further understand the functional relevance of this observation, we studied genetic interactions occurring in strains ablated for both replication proteins. We first monitored bulk DNA replication in synchronized cell cultures and observed that depletion of Fen1 suppressed the replication defect characteristic of *ubp10* mutants, with *ubp10 fen1* cells showing replication dynamics virtually indistinguishable from that of *fen1* single mutants (Figure 2A). We then tested fork transitions upon replication stress induction by dNTP pool depletion, in particular by examining the accumulation of anomalous small Ys in *ubp10*Δ cells^37^. We found that replication intermediates accumulating in *ubp10 fen1* double mutants are very similar to those of *fen1* single mutant cells and clearly differ from those of *ubp10*Δ cells, lacking the characteristic small Y accumulation (Figure 2B), indicating that FEN1 deletion prevents anomalous nascent strand transitions at stalled forks caused by Ubp10 absence. The functional nature of the genetic interaction was further confirmed by testing thermosensitivity and resistance to chronic exposure to HU, where we found that *fen1Δ ubp10*Δ cells phenocopy single *fen1*Δ mutants (Figure 2C), indicating that Ubp10 and Fen1 support viability through a single genetically-related pathway and that Fen1 underlies *ubp10*Δ sensitivity to replication stress. Finally, chromatin fractionation assays in cells synchronously traversing S phase failed to show differences in Fen1 protein accumulation on chromatin in wild-type and *ubp10* defective cells (Figure 2D), suggesting that Ubp10 does not markedly influence Fen1 chromatin association during OFM. Taken together, these analyses suggested that absence of the Fen1 endonuclease, required for Okazaki fragment flap-processing, bypasses a yet undefined replication related Ubp10-dependent function.

**Figure 2.**
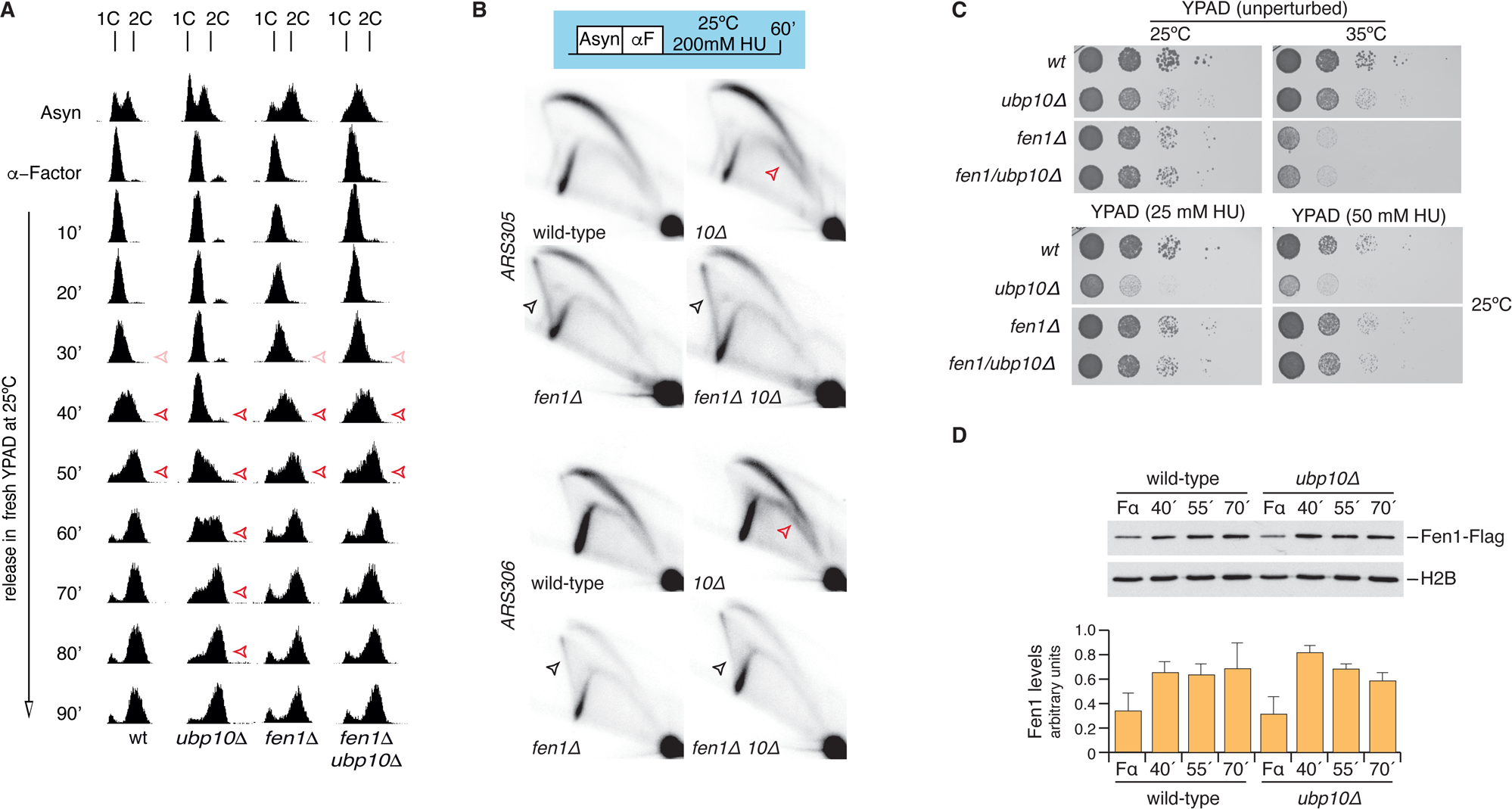
Flap-endonuclease *FEN1/RAD27* is epistatic to PCNA-DUB *UBP10* in the Okazaki fragment maturation pathway. **A.** S phase progression analysis of cells depleted for Fen1 and Ubp10. Wild-type (wt), *ubp10*Δ, *fen1*Δ and *fen1*Δ *ubp10*Δ exponentially growing were synchronized in G1 with α-factor and release in fresh (rich) media. DNA content by FACS analysis of samples taken at indicated intervals is shown. Red arrows indicate approximated duration of bulk DNA replication for each strain. Note that *fen1*Δ *ubp10*Δ cells behave like single *fen1*Δ mutants. **B.** Cells described in A were synchronized with α-Factor and released in the presence of 200 mM HU at 25°C. Samples were taken 60 minutes of treatment and processed for 2D-gel analysis of replication intermediates. Membranes were hybridized consecutively with probes spanning *ARS305* and *ARS306* replication origins. Note that *fen1*Δ *ubp10*Δ cells accumulated X-shaped replication intermediates comparably to *fen1*Δ single mutant cells and in clear contrast to abnormal small Ys intermediates (red arrows) observed in *ubp10*Δ cells (described in Álvarez et al., 2019). **C.** Ten-fold dilutions of the strains indicated in A incubated in YPAD at 25°C or 35°C in the absence (unperturbed) or in the chronic presence of HU (25mM or 50 mM as indicated). Data presented in A, B and C indicates that *FEN1* is epistatic to *UBP10*. **D.** S phase chromatin association of Fen1 in wild-type and *ubp10*Δ cells expressing Fen1-Flag tagged protein. Exponentially growing cultures of wild-type and *ubp10*Δ cells were synchronized with α-factor and released in fresh media to test S phase chromatin association of Flap-endonuclease Fen1/Rad27. Samples were taken at indicated intervals; chromatin-enriched fractions were prepared and electrophoresed in SDS-PAGE gels. Blots were incubated with α-Flag (to detect Fen1-Flag) or α-H2B antibodies. Blots from a representative experiment are shown. Data in the graph represent the average of three biological replicates (and is expressed as means ±SD in triplicate) (p = 0.3250, two-way ANOVA test). This evidence suggests that chromatin association of Fen1 is not affected by depletion of Ubp10.

### Ubp10 promotes Okazaki fragment ligation

In order to explore the role of the Ubp10 in lagging strand synthesis, we analysed chromatin binding of key OFM proteins during unperturbed S phase upon Ubp10 depletion. We show a control assay displaying a chromatin fractionation experiment in full (Supplementary Figure 2), which allows analyzing the partitioning of proteins of interest into chromatin-free and chromatin-bound fractions. For clarity, we only show chromatin-bound fractions in further fractionation assays (including Figure 2D), unless otherwise stated. PCNA is an abundant protein, detectable throughout the cell cycle, that temporarily associates to chromatin during in S phase, in discrete but detectable amounts, in cycles of loading and unloading as the process of DNA synthesis requires^1,8,26^. Indeed, we observed that PCNA is readily detectable in WCE and chromatin-free fractions, and to a lesser extent on chromatin-bound fractions of cells synchronously undergoing genome replication (Supplementary Figure 2). Therefore, during replication, a major fraction of PCNA is detectable in soluble forms while a small subset associates to chromatin with a quite reproducible periodicity.

In *S. cerevisiae, CDC9* encodes the DNA ligase I, an essential enzyme that seals Okazaki fragments during DNA replication^49,50^. Cdc9 has a human homolog, *LIG1*, also regulated by PCNA during the sealing of nicked DNA at lagging strands^51^. Remarkably, human *LIG1* complements yeast *cdc9* temperature-sensitive mutants at the restrictive temperature^52^. Having observed that in budding yeast PCNA-DUB Ubp10 physically interacts with Cdc9 during S phase, we were interested in understanding a possible functional interaction among them.

We studied genetic interactions of *UBP10* with a conditional allele of the DNA ligase I by testing the viability of *cdc9^ts^* and *ubp10*Δ double mutant cells. In our study we used a *cdc9-7* allele (from the National BioResource Project, NBRP Japan) a W303 derivative strain that we characterized and sequenced to find that it is synonymous to the *cdc9-1* allele^53^. We found that depletion of Ubp10 aggravates dramatically the thermosensitivity of strains carrying this *CDC9* temperature-sensitive mutant allele (Figure 3A). We further observed - by chromatin fractionation assays - that abrogation of *UBP10* function in the *cdc9-7 ts* mutant lead to the accumulation of PCNA on chromatin for a longer period of time than in *cdc9-7* controls, which also accumulated PCNA on chromatin when compared to wild-type cells during S phase even under permissive conditions (25°C) (Figure 3B). These observations indicate that the *cdc9-7* allele is - to some extend - defective in nick ligation at 25°C and clearly suggest that all its defects are aggravated by Ubp10 ablation. However, *cdc9-7* mutation did not rescue the characteristic slow bulk DNA replication phenotype of *ubp10*Δ mutant (Supplementary Figure 3A), suggesting that OF ligation is unrelated to the *ubp10*Δ replication defect. We also noticed that deletion of *UBP10* in *cdc9-7* led to an accumulation of unusual replication intermediates by 2D-gel analysis of HU-treated cells equivalent to those observed in *ubp10*Δ single mutants (Supplementary Figure 3B).

**Figure 3.**
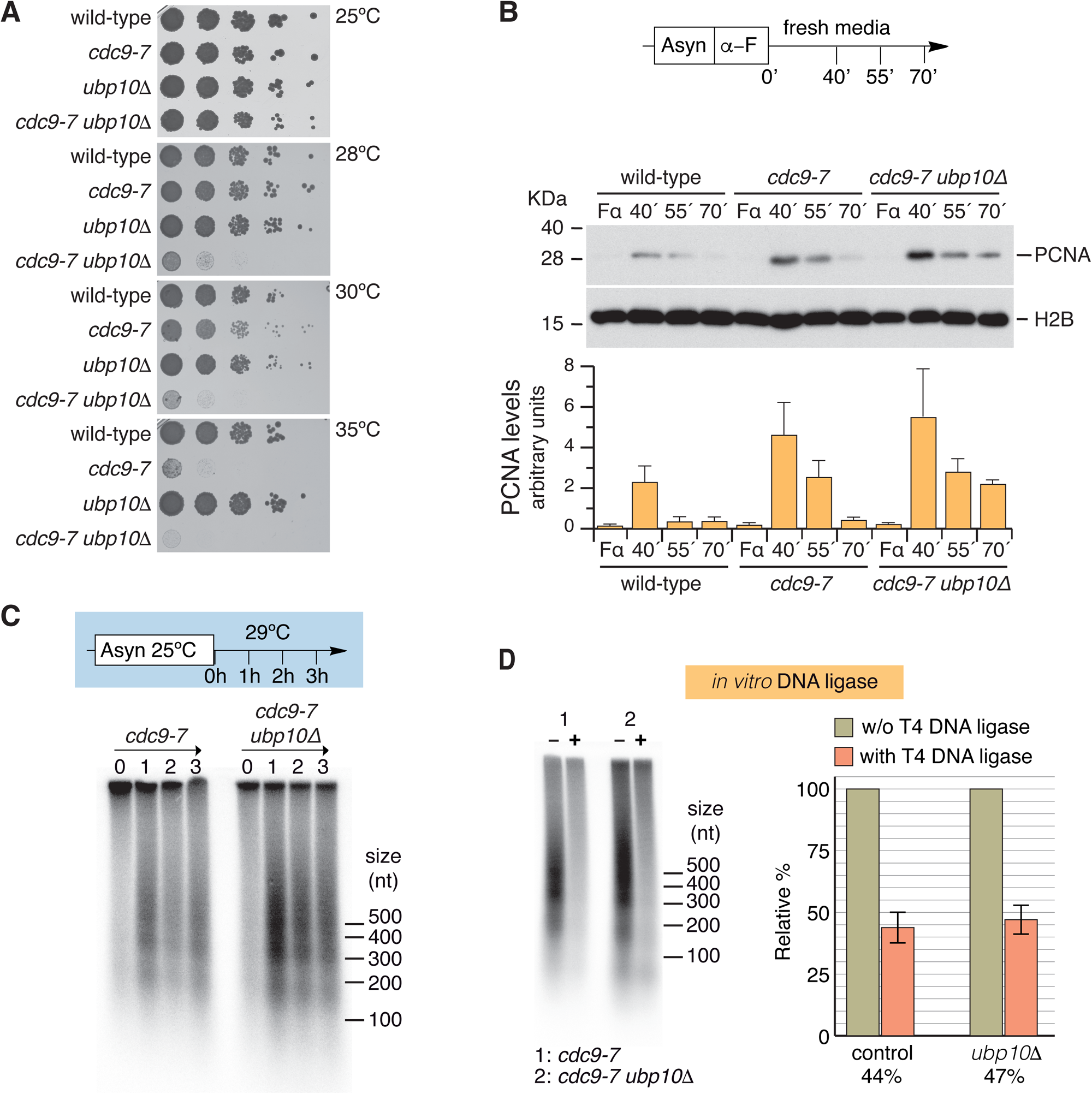
Abrogation of Ubp10 leads to accumulation of unligated Okazaki fragments. Depletion of Ubp10 increases the thermosensitivity and PCNA accumulation defects of the *cdc9-7* ts allele of DNA ligase I. **A**. Ten-fold dilutions of equal number of cells of the indicated strains were spotted in Petri dishes and incubated either at 25°C, 28°C, 30°C, or 35°C for 60 hours. **B**. S phase chromatin association of PCNA in wild-type, *cdc9-7,* and *cdc9-7 ubp10*Δ cells. Exponentially growing cultures of wild-type, *cdc9-7,* and *cdc9-7 ubp10*Δ cells were synchronized in G1 with α-factor pheromone and released in fresh media to test S phase chromatin association of PCNA. Samples were taken at indicated intervals; chromatin-enriched fractions were prepared and electrophoresed in SDS-PAGE gels. Blots were incubated with α-PCNA or α-H2B antibodies. A blot from a representative experiment is shown. Data in the graph represent the average of three biological replicates (and is expressed as means ±SD in triplicate) (wild-type vs *cdc9-7* p = 0.011; *cdc9-7* vs *cdc9-7 ubp10*Δ p = 0.4372, two-way ANOVA test). **C**. *UBP10* mutants show a strong detectable lagging-strand replication phenotype that leads to the accumulation of unligated Okazaki fragments (OF). Exponentially growing cultures of *cdc9-7* and *cdc9-7 ubp10*Δ cells incubated at 25°C were shifted to 29°C. Aliquot samples were taken at the indicated intervals. Purified total genomic DNA was labelled with exonuclease-deficient DNA polymerase I (Klenow) fragment and α-^32^P-dCTP, separated by agarose denaturing electrophoresis, and visualized using a Phosphor Imager. A representative experiment of two biological replicates is shown. Note that *cdc9-7* ts cells transiently accumulate OFs and that depletion of Ubp10 results in further Okazaki fragments accumulation both in quantity and duration. **D**. *In vitro* analysis of *ubp10*Δ cumulative OFs reveals ligatable nick DNA. OFs were obtained from *cdc9-7* and *cdc9-7 ubp10*Δ cells as in C. Where indicated (+) genomic DNA was treated *in vitro* with T4 DNA ligase before labelling with Klenow and α-^32^P-dCTP for quantifying the proportion of DNA fragments (OF) ready for nick ligation. As shown, DNA samples from *cdc9-7 ubp10*Δ cells were as efficiently ligated as *cdc9-7* controls. A representative experiment of three replicates is shown. Values of means of these three replicates ± SD are plotted.

Okazaki fragments can be detected *in vivo* upon DNA ligase I *CDC9* depletion or using conditional alleles of the ligase, which result in the accumulation of nicked DNA^49,50^. To test a potential accumulation of Okazaki fragments in *cdc9-7 ubp10*Δ cells and, particularly, after having observed that *cdc9-7* cells grow poorly at 29°C when combined with deletion of *UBP10*, we set up cultures of exponentially growing cells incubated at 25°C to then shifted them to 29°C. Okazaki fragments were end-labelled in samples taken at one-hour intervals, separated by denaturing agarose gel electrophoresis, transferred to a nitrocellulose membrane, and visualized with a Phosphor Imager. We detected a transitory accumulation of Okazaki fragments in *cdc9-7* cells upon sifting from 25° to 29°C degrees cultures of cells growing exponentially. After end-labelling of DNA and denaturing electrophoresis, we observed the characteristic banding pattern of short and heterogenous nicked DNA that results from defects in the ligation at lagging strands^50^ (Figure 3C). Remarkably, we also found that *cdc9-7 ubp10*Δ cells accumulated OFs more abundantly than *cdc9-7* control cells (Figure 3C). We did not detect this characteristic banding pattern of Okazaki fragments in nick ligation competent wild-type cells or single *ubp10*Δ mutants tested. In essence, all this evidence indicates that depletion of Ubp10 leads to a strong lagging-strand replication defect phenotype. However, it is important to understand whether this strong phenotype is the cause or the consequence of the slow progression through S phase that characterizes the genetic depletion of Ubp10.

Cells with a defective Flap-endonuclease Fen1/Rad27 accumulate unprocessed Okazaki fragments with poorly ligatable ends due to the accumulation of flaps with different sizes^54^ (and our unpublished observations). Furthermore, impairment of PCNA unloader Elg1 function leads to the accretion of extended Okazaki fragments likely reflecting defective post-replicative nucleosome reposition^55^. However, *in vivo* depletion of Cdc9 DNA ligase I causes the accumulation of nicks that are, therefore, *in vitro* ligatable with no obvious defects in coupling with chromatin assembly^50,54^.

To characterize the molecular nature of the Okazaki fragments accumulating in *cdc9-7 ubp10*Δ double mutants, we examined their size by gel electrophoresis and tested the extent to which the isolated DNA fragments are competent for ligation after purification of total DNA (Figure 3D). We observed that *cdc9-7 ubp10*Δ cells accumulate Okazaki fragments of normal length (Figure 3C). In addition, these are *in vitro* ligated by T4 DNA (Figure 3D), with an efficiency equivalent to ligase deficient controls^50,54^. These results indicate that OFs accumulating upon Ubp10 ablation have DNA ends equivalent to those accumulated in *cdc9ts* mutants and suggest that OF accumulation is due to defects in the last steps of lagging strand maturation. The evidence shown so far suggests a role for Ubp10 in the timely maturation of Okazaki fragments, either in the regulation of nick ligation or in promoting chromatin disassociation of PCNA.

### Ubp10 is required for timely ligase association to replicating chromatin

PCNA accumulates on replicating chromatin in the absence of Cdc9-mediated Okazaki fragment ligation^21^. We reasoned that the PCNA accretion observed in Cdc9 Ubp10 doubly depleted cells might be the consequence of defects either in the DNA ligase activity or in the chromatin abundance of Cdc9. Therefore, we next tested a heterologous DNA ligase that can complement Cdc9 depletion^56^, and found that *ubp10*Δ cells overexpressing *Chlorella* virus DNA ligase exhibit a delay in S phase progression similar to that of controls (Supplementary Figure 4A), strengthening the conclusion that *ubp10*Δ deficiency is not related with defects in DNA ligase activity.

We then monitored Cdc9 chromatin association in synchronously replicating cells and found that Ubp10 depleted cells show transiently reduced ligase levels compared to wild type cells (Figure 4), a decrease particularly marked at early time points after G1 release, coinciding with the slow progression of bulk DNA replication and in an inverse correlation with PCNA abundance (Supplementary Figure 4C). Therefore, the accumulation or permanence of PCNA on chromatin in *ubp10*Δ mutants might be the consequence of a slow recruitment of DNA ligase Cdc9 on chromatin. To further understand whether the poor accretion of Cdc9 on chromatin was the cause or a consequence of the slow S phase phenotype of Ubp10 depleted cells, we tested *ubp10*Δ cells overexpressing *CDC9*, by means of a duplicated galactose-inducible allele, and found that high levels of chromatin-bound Cdc9 did not rescue the characteristic cell cycle defect, lengthy S phase, of *ubp10* mutants nor rescued the accumulation of PCNA (Supplementary Figures 4B and 4C).

**Figure 4.**
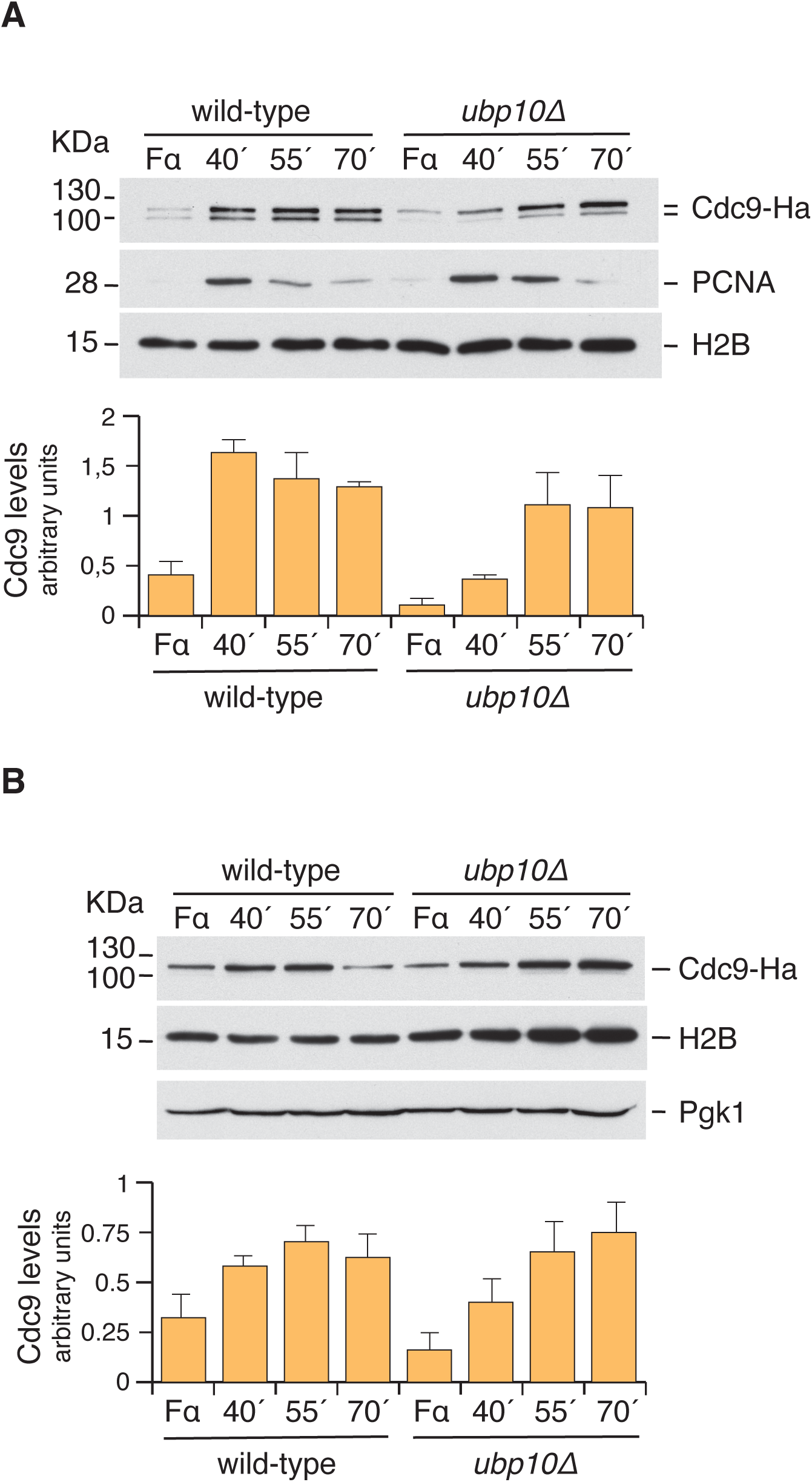
Steady reduction of chromatin associated Cdc9 during replication in *UBP10* defective cells. **A.** S phase chromatin association of Cdc9 in wild-type and *ubp10*Δ cells. Exponentially growing cultures of wild-type and *ubp10*Δ cells were synchronized with α-factor and released in fresh YPAD media to test S phase chromatin association of DNA ligase I Cdc9. Samples were taken at indicated intervals; chromatin-enriched fractions were prepared and electrophoresed in SDS-PAGE gels. Blots were incubated with α-Ha (to detect Cdc9-Ha), α -PCNA or α-H2B antibodies. A blot from a representative experiment is shown. Data in the graph represent the average of three biological replicates (and is expressed as means ±SD in triplicate) (p = 0.0002, two-way ANOVA test). **B.** Whole cell extract (WCE) aliquots from A were blotted to test Cdc9-Ha protein amounts. Again, a blot from a representative experiment is shown. Data in the graph represent the average of three biological replicates (and is expressed as means ±SD in triplicate) (p = 0.1408, two-way ANOVA test).

### PCNA disassembly-prone mutants *pol30^R14E^* and *pol30^D150E^* revert *ubp10*Δ-associated replication defects

We reasoned that the excessive abundance of PCNA on chromatin in Ubp10 depleted cells might be related to defective unloading of the sliding clamp. If *ubp10*Δ phenotypes are related with the excessive retention of PCNA on chromatin, two predictions can be made. Firstly, cells depleted for Ubp10 should accumulate PCNA on chromatin during S phase. Secondly, reduction the amount of PCNA bound to chromatin would rescue *ubp10* replication defects. Regarding this second prediction, there are a number of *POL30* alleles that conform unstable PCNA homotrimers^57^. In particular, *pol30^R14E^* and *pol30^D150E^* alleles are described as PCNA trimer-disassembly-prone mutants due to their instability when bound to chromatin^21,57^. Hence, if the reason underlaying *ubp10* defects is related to excessive PCNA on chromatin, the potential suppression by these disassembly-prone PCNA mutants might be easier to observe in a *cdc9-7 ubp10*Δ background due to the greater thermosensitivity as compared to single *cdc9-7* cells. In fact, the growth defect of *cdc9-7 ubp10*Δ is suppressed by *pol30^R14E^* and *pol30^D150E^* mutant alleles, while they did not rescue *cdc9-7* thermosensitivity (Figure 5A). Furthermore, these two-point mutant alleles of PCNA also rescue abnormal replication intermediates accumulated in *cdc9-7 ubp10*Δ cells arrested in HU (Figure 5B). In fact, these *POL30* alleles rescue all tested Ubp10-depletion associated S phase defects, including those related with bulk genomic DNA replication in a *CDC9* wild-type-like strain (Figure 5C), implying that the slow S phase observed in Ubp10 depleted cells is a consequence of PCNA accumulation on replicating chromatin. Defective replication intermediate accumulation in HU-treated *UBP10* deleted cells is also efficiently suppressed by either *pol30^R14E^* or *pol30^D150E^* (Figure 5D). This evidence indicates that the replication phenotypes caused by Ubp10 ablation are a consequence of a defective PCNA unloading mechanism during S phase.

**Figure 5.**
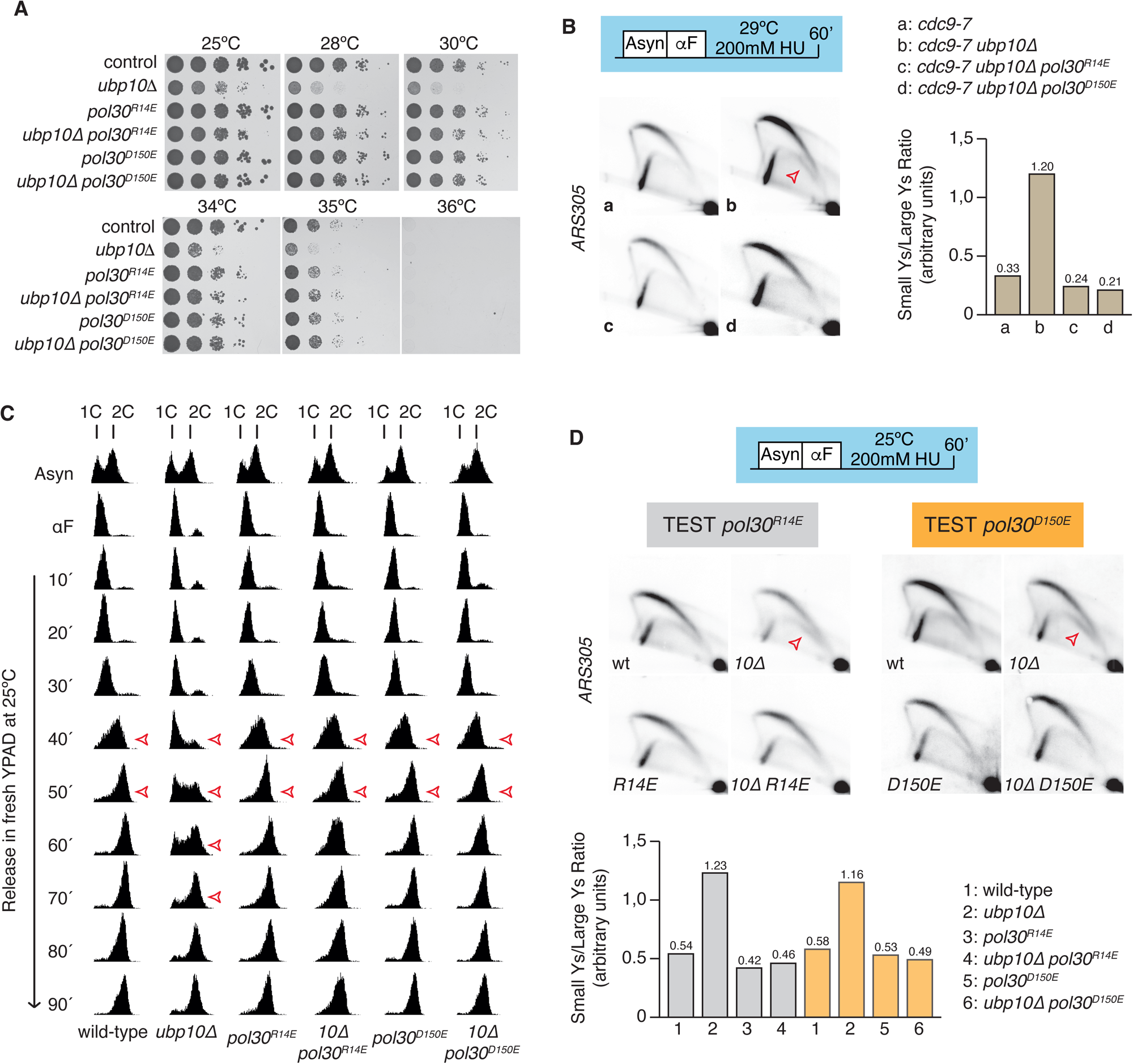
PCNA disassembly-prone mutants *pol30^R14E^* and *pol30^D150E^* revert *ubp10*Δ-associated replication defects. **A.** *pol30^R14E^* and *pol30^D150E^* alleles rescue *ubp10*Δ-associated defects in *cdc9-7 ubp10*Δ. Ten-fold dilutions of the *cdc9-7* indicated strains incubated in YPAD at different temperatures for 60 hours. Note that the increased lethality of *cdc9-7 ubp10*Δ is suppressed by *pol30* mutant alleles that, in turn, do not rescue *cdc9-7* thermosensitivity. **B.** 2D-gel analysis of cells synchronized in early S phase with the ribonucleotide reductase inhibitor HU at 29°C. Indicated *cdc9-7* strains were grown to exponential phase at 25°C, synchronized in G1 with α-factor, released in fresh media with 200 mM HU at 29°C for 60 additional minutes. The membrane was hybridized to a probe spanning *ARS305* early replication origin. Open red arrow points small Ys intermediates. Under these conditions, *cdc9-7 ubp10*Δ defects were suppressed by PCNA disassembly-prone mutants. Histogram plots of small/large Y-shaped replication intermediates ratios in *cdc9-7, cdc9-7 ubp10*Δ, *cdc9-7 ubp10*Δ *pol30^R14E^* and *cdc9-7 ubp10*Δ *pol30^D150E^* mutants are shown. **C.** DNA replication progression defects in *ubp10*Δ cells are abrogated by *pol30^R14E^* and *pol30^D150E^* alleles. DNA content analysis of wild type, *ubp10*Δ, *pol30^R14E^*, *ubp10*Δ *pol30^R14E^*, *pol30^D150E^* and *ubp10*Δ *pol30^D150E^* strains. Cells of the indicated strains (all *cdc9td* with *CDC9* ON) were synchronized with α-factor and released in fresh YPAD at 25°C. The progression of the bulk genome replication was monitored at the indicated time points. Open red arrows indicate approximate S phase duration in every strain where *pol30^R14E^* and *pol30^D150E^* suppression of the replication defect of *ubp10*Δ cells can be observed. **D.** 2D-gel analysis of replication intermediates in cells synchronized in early S phase with HU. Indicated strains grown to exponential phase at 25°C were pre-synchronized in G1 with α-factor, released in fresh media with 200 mM HU an incubated at the same temperature for one additional hour. Membranes were hybridized to a probe spanning *ARS305* early replication origin. Open red arrows indicate small Ys intermediates in blots. Histogram plots of small/large Y-shaped replication intermediates ratios in wild type and *ubp10*Δ, *pol30^R14E^, ubp10*Δ *pol30^R14E^*, *pol30^D150E^* and *ubp10*Δ *pol30^D150E^* mutants are shown.

*UBP10* mutants accumulate non-canonical small Y-shaped replication intermediates upon HU-induced fork stalling^37^. The accumulation of these non-canonical small Ys is characterized by a decrease in large Ys and is abated by mutating *RAD52*^37^. We re-examined the accumulation of these small Y-shaped molecules and found that is suppressed in *pol30^R14E^* and *pol30^D150E^* genetic backgrounds (Figures 5B, 5D). These results strongly suggest that small Y-shaped molecules formed as a direct consequence of PCNA accumulation on replicating chromatin (and, also, that these abnormal structures are generated as cells try to repair through a Rad52-dependent TS-like mechanism).

### Disassembly-prone-mutant *pol30^D150E^* suppresses the increased chromatin association of PCNA as well as the OF accumulation in *UBP10* defective cells

As mentioned, if *ubp10* phenotypes are the consequence of a defective unloading of PCNA, cells depleted for Ubp10 should accumulate the sliding clamp on chromatin during S phase, indeed we have observed a chromatin-bound accretion in *ubp10*Δ cells when testing Cdc9 levels (Figure 4A and Supplementary Figure 4C). We next re-evaluated how much and for how long the sliding clamp PCNA is bound to chromatin through a synchronized, otherwise unperturbed, S phase in wild-type and *ubp10*Δ cells. Exponentially growing cultures of either wild-type or *ubp10*Δ strains were synchronized in G1 with α-factor pheromone and release in fresh media. Samples were taken at 0, 40, 55 and 70 minutes after the release and processed for chromatin fractionation assays and DNA content analysis. As expected from previous results (Figure 5C), we found that the slow S phase progression in *ubp10*Δ cells correlates with an increased PCNA accumulation on chromatin (Figure 6A and 6B). In parallel, we also evaluated the disassembly-prone *pol30^D150E^* point-mutant ability to suppress this chromatin-bound PCNA accretion phenotype and found that this PCNA mutant rescued chromatin retention of PCNA of *ubp10*Δ (Figure 6A), as well as the bulk DNA replication defect (Figure 5C and 6B).

**Figure 6.**
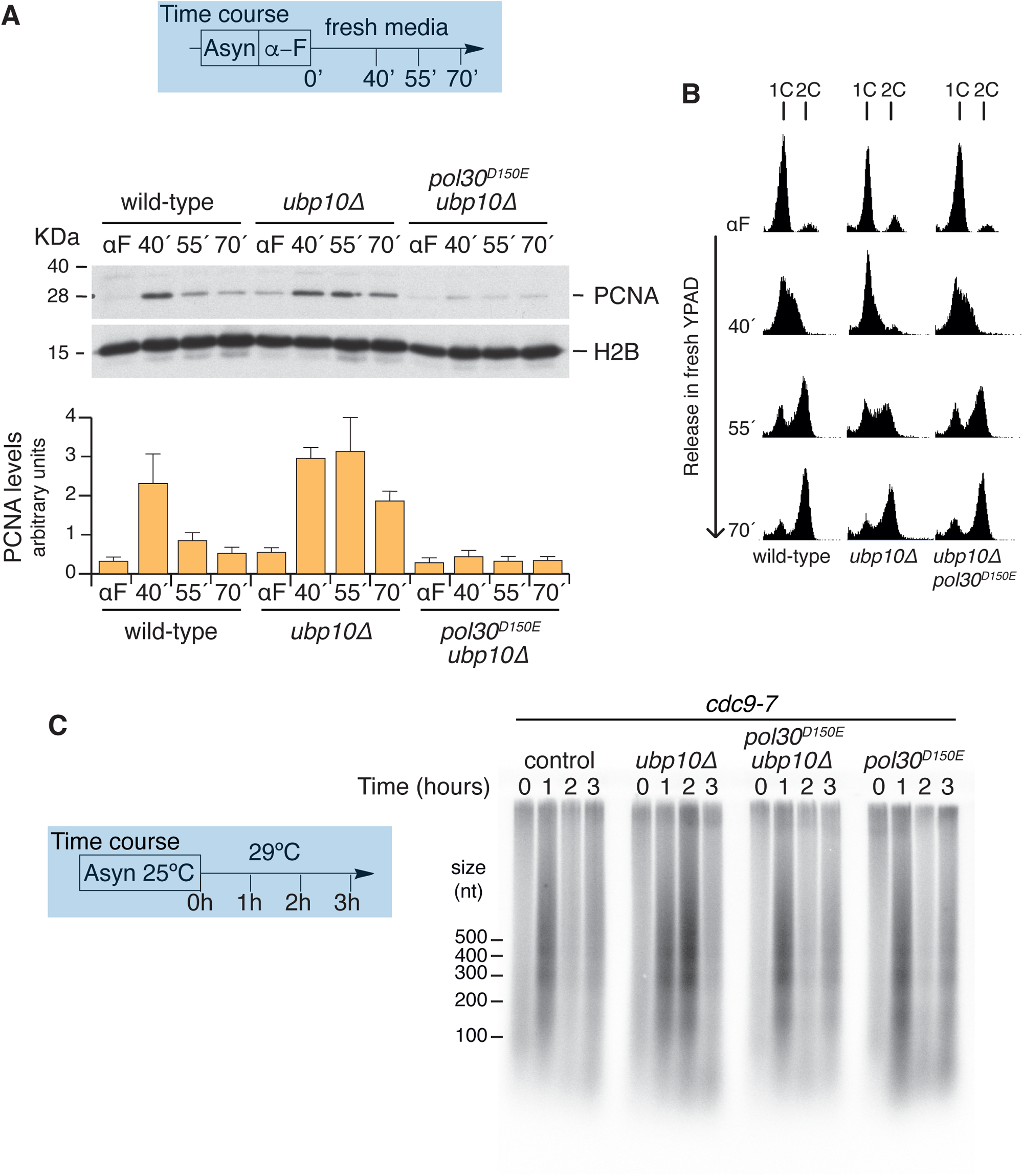
PCNA retention on replicating chromatin underlies Okazaki fragments accumulation in Ubp10 abrogated cells. **A.** PCNA trimer-disassembly-prone mutant *pol30^D150E^* suppresses PCNA retention on chromatin phenotype of Ubp10 depleted cells. Exponentially growing cultures of wild-type, *ubp10*Δ, and *pol30^D150E^ ubp10*Δ cells were synchronized in G1 with α-factor and released in fresh media to test S phase chromatin association of PCNA. Samples were taken at indicated intervals; chromatin-enriched fractions were prepared and electrophoresed in SDS-PAGE gels. Blots were incubated with α-PCNA or α-H2B antibodies. A blot from a representative experiment is shown. Data in the graph represent the average of three biological replicates (and is expressed as means ±SD in triplicate) (wild-type vs *ubp10*Δ p <0.0001; *ubp10*Δ vs *pol30^D150E^ ubp10*Δ p <0.0001, two-way ANOVA test). **B.** Slow S phase progression in PCNA-DUB *UBP10* defective cells is a direct consequence of PCNA accumulation on replicating chromatin. DNA content analysis of wild type, *ubp10*Δ, and *pol30^D150E^ ubp10*Δ cells at the indicated time points (aliquot samples of the experiment in A). **C.** *pol30^D150E^* rescues *ubp10*Δ Okazaki fragments maturation timing defects. Exponentially growing cultures of *cdc9-7*, *cdc9-7 ubp10*Δ, *pol30^D150E^* and *pol30^D150E^ ubp10*Δ cells incubated at 25°C were shifted to 29°C. Aliquot samples were taken at the indicated one-hour intervals. Purified total genomic DNA was labelled with exonuclease-deficient DNA polymerase I, Klenow, fragment and α-^32^P-dCTP, separated by agarose denaturing electrophoresis, and visualized using a Phosphor Imager. A representative experiment of two biological replicates is shown.

Having shown that PCNA accumulation on replicating chromatin underlies not only the slow S phase progression but also the formation of anomalous small Ys in *ubp10* defective cells, we next tested whether the PCNA disassembly-prone *pol30^D150E^* allele would be able to rescue the accumulation of unligated Okazaki fragments that can be evidenced in the *ubp10* mutant using a *cdc9-7* background. It has been shown that *pol30^R14E^* and *pol30^D150E^* PCNA trimer-disassembly-prone mutants alleviate the Okazaki fragment length extension problem described in *elg1*Δ cells^55^, in agreement with PCNA unloading being a key event in maturation of the lagging strand and nucleosome deposition. We predicted that *pol30^D150E^* would mitigate the OF accretion of *cdc9-7 ubp10*Δ cells. To test this hypothesis, exponentially growing *cdc9-7*, *cdc9-7 ubp10*Δ, *pol30^D150E^* and *pol30^D150E^ ubp10*Δ cells incubated at 25°C were shifted to 29°C, and samples were taken at one-hour intervals and processed for OF analysis. We observed that the *pol30^D150E^* point mutant suppresses *ubp10*Δ Okazaki fragments maturation timing defect (Figure 6C), implying that the main problem caused by Ubp10 depletion during lagging strand replication is PCNA retention on chromatin.

### Ubp10 as a key regulator of a PCNA unloading mechanism distinct from Elg1-RLC

Thus far, our results argue that Ubp10 promotes timely PCNA unloading during lagging strand replication. One reasonable hypothesis is that Ubp10 promotes PCNA deubiquitylation to enhance Elg1-mediated PCNA unloading at the final steps of Okazaki fragment maturation. Fully aware of *in vitro* evidence showing that human ATAD5^Elg1^ is able to unload both ubiquitylated and deubiquitylated PCNA forms with similar efficiency^58^, we reasoned that *in vivo*, during unperturbed DNA replication, chromatin PCNA unloading would be enhanced by deubiquitylated forms of the sliding clamp in order to proceed smoothly and timely through lagging strand synthesis.

Yeast Elg1^ATAD5^ is an evolutionary conserved homolog of the replication factor C (RFC) subunit Rfc1^23,25^. It has been shown that while Rfc1-RFC loads PCNA on replicating chromatin, the Elg1-RLC complex has a role in PCNA unloading in a molecular event preceded by Okazaki fragment ligation^18,21^. Elg1 forms an alternative RFC hetero-pentameric complex with all RFC2-5 subunits of RFC. Significantly, this alternative Elg1-RLC complex is important but not essential for DNA replication^24^. Elg1 interacts physically with PCNA and Fen1^Rad27^, has a role in PCNA unloading during Okazaki fragment maturation and is, therefore, important for efficient S phase progression, and likely has a role in proper nucleosome assembly^18,25,55^.

We have observed that, though Elg1 displays many phenotypes related to chromosome stability, depletion of *ELG1* in budding yeast do not cause major replication delays as assayed in synchronous S phase (Supplementary Figure 5A). In fact, bulk DNA replication timing in *ELG1* mutants is equivalent to wild-type replication as tested by FACS DNA content analysis (Supplementary Figure 5A). Moreover, deletion of *ELG1* is viable while deletion of other RFC components is not (in particular RFC1)^24,26^ (Supplementary Figure 5B). However, in 10-fold dilution assays we detected that *elg1*Δ strains show a poor growth rate particularly at high temperatures (Supplementary Figure 5B). A defect exacerbated when single *elg1*Δ mutation is combined with the deletion of Ubp10 (*ubp10*Δ *elg1*Δ) at any tested temperature, as compared to single mutants or wild type cells (Supplementary Figure 5B). This semi-lethality is indicative of a genetic interaction suggestive of a role for both factors in a common event, likely PCNA unloading.

To test chromatin-bound PCNA levels throughout S phase in wild-type, double mutant *elg1Δ ubp10*Δ, and single mutants *elg1*Δ and *ubp10*Δ, mid-log phase cultures of the indicated strains were synchronized in G1 with α-factor and released in fresh media. As in previous experiments, samples were taken at indicated time points and processed for chromatin pellet assays and DNA content analysis (Figure 7). We initially expected *elg1Δ ubp10*Δ double mutants to behave like *elg1*Δ singles in accordance with the hypothesis that Ubp10 might regulate PCNA unloading through Elg1. Unexpectedly, we found that *ubp10*Δ and *elg1*Δ, when combined, are additive regarding PCNA accumulation on replicating chromatin (Figure 7A and 7B), strongly suggesting the existence of two distinct pathways of PCNA unloading. These results support the existence of a chromatin disassociation of PCNA mechanism regulated by Ubp10 separable from the Elg1^ATAD5^-dependent PCNA unloading.

**Figure 7.**
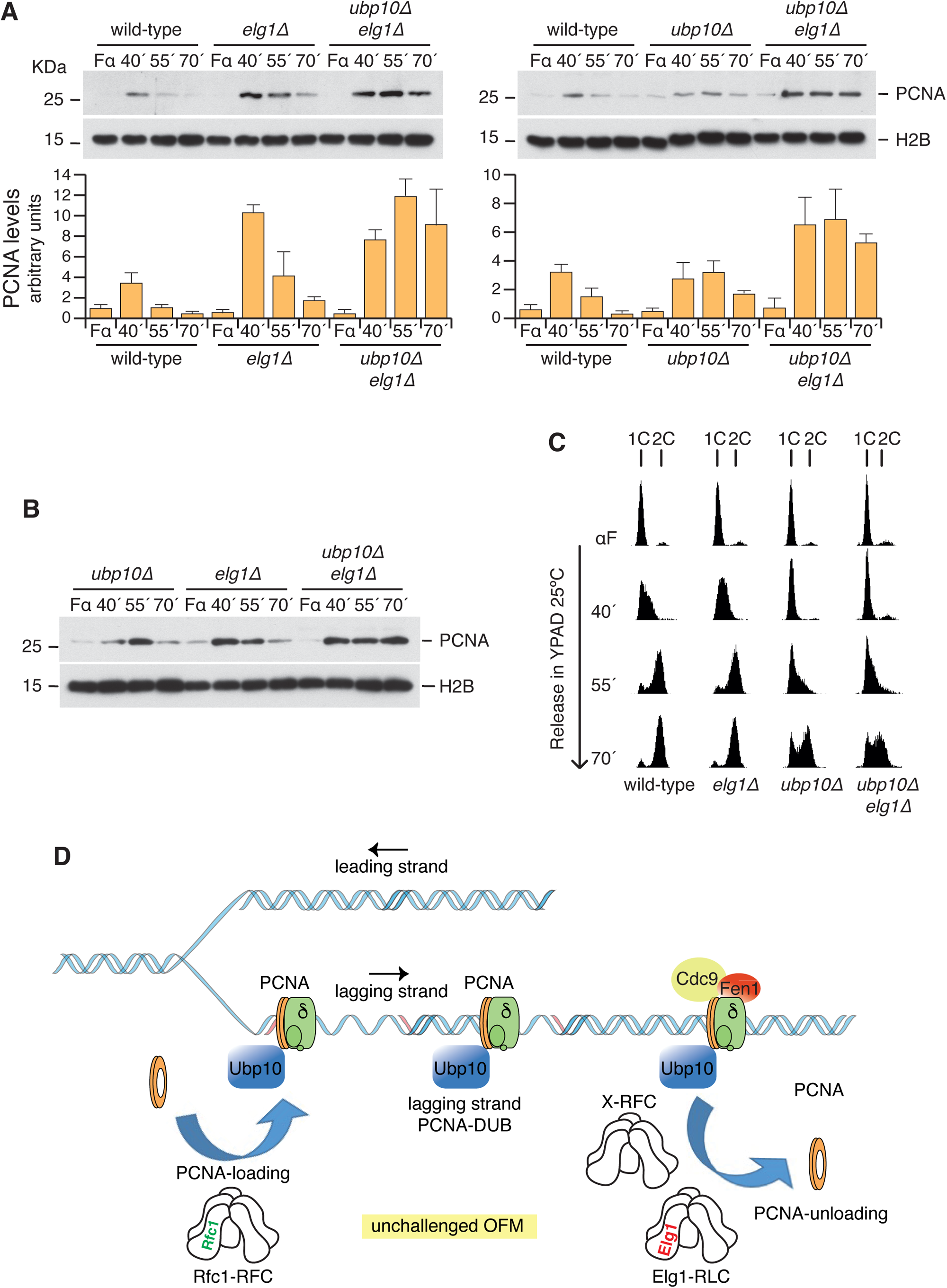
Ubp10 is a key regulator of a PCNA unloading mechanism distinct from Elg1-RLC. **A.** Abrogation of Ubp10 is strongly additive to Elg1 depletion resulting in substantial increase of PCNA bound to chromatin during replication. S phase chromatin association of PCNA in wild-type, *elg1*Δ, *ubp10*Δ and *elg1Δ ubp10*Δ cells. Exponentially growing cultures of the indicated strains were synchronized in G1 with α-factor and released in fresh media to test S phase chromatin association of PCNA and histone H2B. Samples were taken at indicated intervals; chromatin-enriched fractions were prepared and electrophoresed in SDS-PAGE gels. Blots were incubated with α-PCNA or α-H2B antibodies. Blots from representative experiments are shown. Data in the graphs represent the average of three biological replicates (and are expressed as means ±SD in triplicate) (p < 0.0001, two-way ANOVA test). Note that depletion of Elg1 results in transient accumulation of PCNA on chromatin during S phase. **B.** S phase chromatin association of PCNA and histone H2B in *ubp10*Δ, *elg1*Δ and *elg1Δ ubp10*Δ cells. Samples were processed as in A to compare the transient accumulation of PCNA on replicating chromatin in *ubp10*Δ and *elg1*Δ single mutants in the same blot (p < 0.0001, two-way ANOVA test). **C.** S phase progression analysis of wild-type, *elg1*Δ, *ubp10*Δ and *elg1Δ ubp10*Δ cells. Progression of bulk genome replication was monitored at the indicated time points by FACS analysis. **D.** Proposed model for PCNA-DUB Ubp10 during maturation of Okazaki fragments regulating an Elg1-independent PCNA unloading mechanism (see text for details).

Even though the proteomic analysis presented in this work revealed no interaction with the RFC cofactor Elg1, we examined whether the ablation of Ubp10 altered the chromatin binding pattern of Elg1 during a synchronized S phase. By chromatin fractionation assays, we found that depletion of the PCNA-DUB does not alter Elg1 interaction with chromatin (Supplementary Figure 5C). Therefore, we concluded that *UBP10* mutants do not deregulate Elg1 interaction with chromatin during S phase. Significantly, cells depleted for both *UBP10* and *ELG1* do not exacerbate the *ubp10*Δ deletion slow S phase phenotype, though they accumulate PCNA abundantly, far more than individual mutants (Figure 7A and 7B). The fact that depletion of *UBP10* alone has a replication progression defect underpins the importance of the PCNA-DUB Ubp10-dependent mechanism of PCNA unloading during lagging strand synthesis.

## DISCUSSION

Okazaki fragment maturation is a complex, yet well understood, process in the synthesis of the lagging strand during DNA replication. Here, we unveil a role of the ubiquitin protease Ubp10 in the latest steps of maturation of Okazaki fragments in the model yeast *S. cerevisiae*. In yeast, the Ubp10 enzyme have been functionally related to RNA polymerase I through the stabilization of the Rpa190 subunit^41^. Ubp10 also cooperates with the FACT complex in the maturation of nucleosomes^48^. Furthermore, Ubp10 is involved in the reversal of histone H2B^K123^ ubiquitylation^38–40^. Significantly for this report, a key role of this desubiquitylase is to counteract futile bypass events at replication forks acting as a PCNA^K164^–DUB^37^. PCNA deubiquitylation is a requirement conserved throughout evolution as evidence for ScUbp10, SpUbp16 and HsUsp1 shows^36,43^. With this background in mind, the aim of this work was to understand the functional meaning of the interaction of the PCNA-DUB Ubp10 with proteins involved in Okazaki fragment synthesis and maturation, physical interaction described earlier for the Flap endonuclease Fen1^Rad27,37^. In this study we have presented ample evidence suggesting that this DUB regulates the dissociation of the sliding clamp PCNA from chromatin and that, by doing so, ensures proper maturation of the lagging strand.

One recent observation, Fen1^Rad27^-Ubp10 binding, led us to the study of Ubp10’s S phase proteome. The analysis confirmed previous data regarding Ubp10 biology as the RNA polymerase I complex subunits Rpa190, Rpa34, Rpa43, Rpa49 and Rpa135 were among proteins trap with Ubp10. Spt16 and Pob3 FACT subunits were also found to bind Ubp10-GFP. Our approach confirmed Fen1-Ubp10 and PCNA-Ubp10 interactions and identified major components of synthesis and maturation of the lagging strand as feasible interactors of the PCNA-DUB. The proteomic studies were made in crosslinked protein samples from cells synchronized in S phase. We confirmed each observed interaction by individually testing Ubp10 ability to form a complex with each OFM complex component of interest in tagged strains during S phase.

A relevant point for this work is understanding the nature of the cell cycle defect of Ubp10 depleted cells, a defect we believed is poorly understood^36,37,41,42^. Growth and cell cycle defects are separable^37,42^. In our studies, we did not observe a G1 delay defect^36,37^. Further, the timing of entry into S phase is close to that of the wild type, with equivalent timing in ARSs activation^37^. *ubp10*Δ characterization indicates that the defective cell cycle is a consequence of the slowdown in DNA replication progression. We also found here that this defect is based on the accretion of chromatin-bound PCNA during S phase and that, consistent with this evidence, it is efficiently suppressed by PCNA-disassembly-prone *pol30^R14E^* and *pol30^D150E^* mutant alleles (see below).

Fission yeast cells increase the amount of chromatin-associated PCNA when the K164 of this sliding clamp is ubiquitylated^44^. Based on their observations, Daigaku and coworkers proposed that in *S. pombe* an increase in Ub-PCNA^K164^ works to expand the time for PCNA-Pol δ binding to chromatin to allow the completion of OFs. Our findings in *S. cerevisae* are consistent with a scenario where ubiquitylation of PCNA is a DNA retention signal for the sliding clamp at lagging strands in unperturbed replication. In a cause-and-effect link (further discussed below), the increase in chromatin-bound PCNA likely slows down progression through S phase in budding yeast. This is consistent with previous evidence in *S. pombe* cells where depletion of PCNA-DUBs leads to a cell cycle delay phenotype suppressed by abrogation of the PCNA-ubiquitin-ligase Rhp18^43^.

Of particular interest for our work were both the study of the cell cycle defect of Ubp10 depleted cells and the analysis of the genetic interactions that arise from individual mutants in the OFM pathway when combined with *ubp10*Δ. One of these analyses has been made in a Cdc9 defective background. In this analysis, not only we found a semi-synthetic lethality among *cdc9-7* and *ubp10*Δ but also created a tool and found a temperature for the OF accumulation tests. Indeed, *cdc9-7 ubp10*Δ results suggest that ablation of the PCNA-DUB Ubp10 leads to a strong lagging-strand replication defect phenotype, possibly owing to defects in Okazaki fragment ligation. On the other hand, we also show that *ubp10*Δ phenotype is unrelated to Cdc9 presence or activity. Therefore, we surmise that *ubp10*Δ cells have a genuine defect in the maturation of Okazaki fragments related to PCNA unloading.

We have performed a series of experiments to measure the length and molecular nature of the abundant OF accumulated in *UBP10* mutant cells and, in summary, all observations indicate that only abundance is affected. Regarding the length and relative accumulation of chromatin repeats (that reflect nucleosome repeats), the detected OF in *cdc9-7 ubp10*Δ cells are normal. Moreover, we show here that Ubp10 accumulated OF are in vitro ligatable to the same extent as controls (*cdc9-7 ubp10*Δ versus *cdc9-7*), and controls are in accordance with published data^54^ again indicating that OFs are conventional nicked DNA. Since it has been demonstrated that lagging-strand synthesis in budding yeast is coupled with chromatin assembly on newly synthesized DNA^50^ we deduce as well that chromatin assembly is not affected in Ubp10 depleted cells.

Smith and Whitehouse have shown that *rad9*Δ and *tof1*Δ checkpoint mutants when combined with a *cdc9^td^*-degron allele accumulate abundant but normal length Okazaki fragments when Cdc9^td^ is proteolyzed^50^. This observation is particularly strong, in terms of OF abundance, for *tof1*Δ mutants and is relevant for our work given the similarity with *ubp10*Δ data presented here. There is not such a strong accumulation in *cdc9^td^ rad9*Δ case^50^. Tof1, named after topoisomerase I-interacting factor, is pertinent to this work because is a subunit of the Csm3-Mrc1-Tof1 replication pausing-mediator complex functionally associated with DNA replication forks^59–62^. On the other hand, the N-terminus of FACT-subunit Spt16 interacts with Tof1 to ensure chromatin replication in vitro^63^, supporting the hypothesis that FACT is recruited to replication forks by the Tof1-fork replication complex for parental nucleosomes removal. Given that the FACT complex interacts with Ubp10 likely to integrate (transcription and) DNA replication with nucleosome assembly^48^, a complex Tof1-FACT-Ubp10 connection emerges likely involved in robust DNA replication progression.

As mentioned before, an important conclusion here is that *ubp10*Δ S phase defects do not involve Cdc9 function. However, we observed that *ubp10*Δ cell cycle deficiency was assumably a consequence of increased residence time of PCNA during S phase and was directly related with a defective unloading of the sliding clamp in Ubp10 depleted cells. Accordingly, disassembly-prone mutant alleles of *POL30*, *pol30^R14E^* and *pol30^D150E^*, rescue *ubp10*Δ mutant defects. A corollary of these two remarkable observations is that during unperturbed S phase PCNA unloading may occur simultaneously with or even before Cdc9-mediated nick ligation and this may be in contradiction with evidence published regarding Elg1 in the PCNA unloading subject^21^. However, this is an outstanding preliminary observation; therefore, further work would be required to understand this conundrum.

Besides the slowdown in replication progression, the ablation of Ubp10 is characterized by the accumulation of non-canonical replication intermediates in HU-treated cells detected as small Ys by 2D-gel analysis. These intermediates are observed upon Ubp10-DUB depletion, suggesting that they are normally suppressed by PCNA deubiquitylation^37^. Canonical small Ys reflect passive replication by forks arising outside the probed fragment. We suggest that accumulation of small Ys may reflect also pathological features of lagging strand-associated replication fork defects (nick DNA that may generate breakage structures). Special consideration should be given to replication intermediates when testing drug-treated cells because fork progression is limited in HU, as elongation of DNA synthesis from ARSs is slow^64^, thus, all replication intermediates detected under our experimental conditions by 2D gel analysis belong to the closest origin of replication for any given restriction fragment tested. Consistent with the idea that non-canonical/small Ys might be pathological replication structures, both *fen1^rad27^* and *cdc9-7* mutants accumulate small Ys (comparable to *top1 top2* double mutants^65^) compatible with increased accumulation of nicked DNA, therefore, fragile molecules.

The suppression of the accumulation of these non-canonical molecules in HU-treated cells by *POL30* disassembly-prone mutant alleles, *pol30^R14E^* and *pol30^D150E^*, combined with the suppression of the slow bulk DNA replication, show that *ubp10*Δ mutation have an effect throughout the entire yeast genome. All this evidence indicates that this Ubp10 deubiquitylase plays a significant genome-wide role during the S phase of every unperturbed cell cycle.

PCNA trimer-disassembly-prone mutants *pol30^R14E^* and *pol30^D150E^* alleviate the Okazaki fragment length extension problem described in *elg1*Δ cells^55^. Further, combining *pol30^R14E^* with *elg1*Δ largely rescued the elevated mutation rate of the single *elg1* mutant^57^. Although *UBP10* mutant cells accumulate OFs, they do not show an elevated mutation rate, in fact their mutation rate is similar to that of wild-type cells^36,43^. Both, *pol30^D150E^* and *pol30^R14E^*, PCNA mutant alleles are excellent extragenic suppressors of *UBP10* deletion. It has been described that due to their disassembly-prone nature of the homo-trimeric PCNA ring, both *POL30* alleles accumulate low levels of PCNA on replicating chromatin. However, based in our own observations, *pol30^R14E^* and *pol30^D150E^* point mutant alleles differ between them in the total cellular amount of PCNA levels. In whole cell extracts, PCNA^D150E^ levels mimics those of wild-type cells, while PCNA^R14E^ shows significantly reduced levels as compared to PCNA^wt^. Thus, PCNA^D150E^ behaves like a real prone-disassembly PCNA mutant. Nevertheless, PCNA^R14E^ mirror PCNA^D150E^ low levels of PCNA bound to chromatin and, therefore, it is useful and valuable in our analysis.

In summary, disassembly-prone mutants *pol30^R14E^* and *pol30^D150E^* rescue the chromatin retention of PCNA phenotype of *ubp10*Δ (as shown in Figure 6A), as well as cell cycle delay (Figure 5C and 6B) and defective replication intermediate accumulation upon exposure to HU (Figure 5D). Remarkably, the pol*30^D150E^* disassembly-prone mutant also rescues Okazaki fragments accumulation observed in DNA ligase I (*cdc9ts*) when combined with *ubp10*Δ (Figure 6C). In other words, all major S phase phenotypes associated with defective Ubp10 (*ubp10*Δ) are relieved by reversion of accretion of PCNA by two different trimer instability PCNA/*POL30* mutants (*pol30^R14E^* and *pol30^D150E^*)^55,57^. Together, all this evidence indicates that a slow PCNA unloading underlies every replication defect in Ubp10 depleted cells.

Another relevant information here is that Elg1-depleted cells transiently accumulate PCNA on chromatin (Figure 7). This transient nature of PCNA retention on chromatin is, on one hand, consistent with the viability *elg1*Δ deleted strains^23–25^. However, on the other hand, it may also mean that in the absence of Elg1-RLC complex PCNA is steadily unloaded in vivo, as we show here (Figure 7). Although it does not come as a total surprise, this unanticipated observation is coherent with the fact that Elg1 is not essential for DNA replication^23–25^. Nonetheless, as reported, Elg1 may be still important for efficient S phase progression^18^. However, by testing bulk DNA replication in unperturbed conditions at 25°C, we observed no S phase delays in Elg1-depleted cells (in similar conditions where we detected a transient accretion of chromatin-bound PCNA). One strong possibility is that under these experimental circumstances an alternative complex to Elg1-RLC1 is unloading PCNA.

The transient accumulation of chromatin-bound PCNA in *elg1*Δ cells is, therefore, evocative of the existence of an alternative PCNA unloading mechanism during replication. We show here that only when *elg1*Δ is combined with *ubp10*Δ PCNA remain bound onto chromatin. Interestingly, we detected a robust accumulation and a very slow-paced PCNA unloading in *elg1 ubp10* double-mutant cells in synchronized S phase, suggesting the existence of an Ubp10-regulated PCNA unloading mechanism divergent from that of Elg1-RLC. Undeniably, all the evidence suggests that a Ubp10-dependent mechanism underlies the timely removal of PCNA from replicating chromatin during unperturbed S phase. Eventually, PCNA is unloaded in Elg1 and Ubp10 doubly-depleted cells, as chromatin-bound PCNA in G1 synchronized cells remains very low. Perhaps most significantly, S phase progression is slow in this *elg1 ubp10* double mutant. The fact that depletion of Ubp10 has a similar slow S phase phenotype may be indicative of a default mechanism of PCNA unloading regulated by this PCNA-DUB in unperturbed cell cycle.

Kubota et al., predicted Elg1-RLC alternative PCNA unloaders back in 2013^26^. In fact, they suggested three models of PCNA unloading based on published evidence at the time and still valid today. A first model where Elg1-RLC would be the main unloader; a second model where Rfc1-RFC would work as a genome-wide unloader and Elg1-RLC would be the unloader of PCNA at specific genome localizations with emphasis in difficult to replicate sites; and a third model where Elg1-RLC complex would unload SUMOylated PCNA and Rfc1-RFC would unload unmodified PCNA rings to directly recycle them at lagging strands. Our findings are consistent with the last two models yet better support the third one where two different PCNA unloaders complexes would effectively recycle PCNA during the synthesis of lagging strands. We suggest that the most likely alternative PCNA unloader is Rfc1-RFC, and that it may unload PCNA preferentially when deubiquitylated. Rfc1 is an essential subunit of the RFC complex that interacts with Ubp10 and PCNA. Ubp10 would eventually regulate the precise timing of PCNA unloading by Rfc1-RFC after Ub-PCNA^K164^ deubiquitylation. Our working model predicts that PCNA is ubiquitylated at K164 at lagging strands, we hypothesize that such event would be a consequence of the collision of the PCNA-Pol delta complex with the preceding (5’ end) Okazaki Fragment. Prior to Ubp10 action and subsequent PCNA unloading, PCNA ubiquitylation would enhance Pol delta release by a collision-release mechanism already described^66^.

A strong case can be made here that yeast cells coordinate the last steps of synthesis and maturation of Okazaki fragments through a Ubp10-regulated PCNA unloading mechanism distinct from that of Elg1^ATAD5^-RLC (see model in Figure 7D). Our findings are consistent with the hypothesis that *S. cerevisiae* cells ubiquitylate and deubiquitylate PCNA during S phase in a dynamic, yet ordered, manner to ensure normal DNA replication, such that PCNA ubiquitylation is orderly followed by its deubiquitylation to strengthen PCNA unloading at lagging strands in a genome-wide scale, regulating and ensuring the time frame for Okazaki fragments maturation to generate a continuous lagging double-stranded DNA.

## MATERIAL AND METHODS

### Yeast strains, growth conditions and media

All the budding yeast used in this study originate from a *MATa* W303 *RAD5 bar1::LEU2* strain^36^ and are listed in the Resources Table (Supplementary Information). Budding yeast strains were grown in YPAD medium (1% yeast extract, 2% peptone supplemented with 50 μg/ml adenine) containing 2% glucose. For block-and-release experiments, cells were grown in YPAD with 2% glucose at 25°C and synchronized in G1 with α-factor pheromone (40 ng/ml, 2.5 hours). Cells were then collected by centrifugation (3000 rpm, 3 min) and released into fresh media (supplemented with 50 µg/ml of Pronase) in the absence or in the presence of HU (0.2 M, FORMEDIUM). Overexpression experiments with cells grown in YPAD medium with 2% raffinose at 25°C were conducted by adding to the medium 2.5% galactose (to induce) or 2% glucose (to repress).

### General experimental procedures

General experimental procedures of yeast Molecular and Cellular Biology were used as described previously^60,67,68^. Generation of tagged alleles and specific gene deletions was performed as described^69^. Transformation was performed by lithium acetate protocol and transformants were selected by growing in selective mediums. Different selection markers were used (*KANMX6*, *HphMX4*, *NatMX6*, *URA3*, *TRP1*, *HIS3*), as indicated in Resources Table. Constructs were confirmed by PCR and/or sequencing. The presence of tagged proteins was further confirmed by immunoblot. Moreover, strains with tagged alleles were carefully checked for growth rate and sensitivity to HU. No differences with untagged controls were found.

### Flow Cytometry Analysis

For flow cytometry analyses, 10^7^ cells were collected by centrifugation, washed once with water, fixed in 70% ethanol, and processed as described previously^68,70^. Cells were prepared using a modification of the method, by using SYTOX Green (Molecular PROBES) for DNA staining^71,72^. The DNA content of individual cells was measured using a Becton Dickinson Accuri C6 plus FACScan.

### HU sensitivity Assays

Stationary cells were counted and serially diluted in YPAD media. Ten-fold dilutions of equal numbers of cells were plated onto YPAD (2% glucose) media (always supplemented with 50 μg/ml adenine), or YPAD containing HU, incubated at the indicated temperatures for 24, 48, 72 or 120 hours and then scanned.

### Identification of Ubp10 interactors by mass spectrometry

Ubp10-GFP expressing- and wild type control cells were synchronized with α-factor and released into fresh YPAD. 30 min, 40 min, 50 min and 60 min time point samples were fixed with formaldehyde and harvested. Cell pellets were resuspended in Lysis Buffer (50 mM Hepes pH 7.5, 140 mM NaCl, 1 mM EDTA, 1% Tritón-X100, 0.1% Na-deoxycholate) and broken using glass beads in a fast-prep. Chromatin extracts were disrupted by sonication, cleared by centrifugation, and incubated for 4 hours at 4°C with agarose-conjugated GFP-Trap™ beads (Chromotek). Beads were washed once with lysis buffer, once with Wash Buffer (10 mM Tris pH 8, 250 mM LiCl, 1 mM EDTA, 0.5% NP-40, 0.5% Na-deoxycholate) and once with TE. 30 min, 40 min, 50 min and 60 min samples were pulled and resuspended in Laemmli Buffer, and resolved by SDS-PAGE. Regions of interest were excised and digested as previously described^73^. Samples were digested using trypsin and analyzed using mass spectrometry at the Proteomic Biotechnology Unit of the Cancer Research Center (Salamanca, Spain). The mass spectrometry proteomics data have been deposited to the ProteomeXchange Consortium via the PRIDE^74^ partner repository with the dataset identifier PXD048249.

### Total protein extracts for Immunoblotting

Total protein extracts were prepared following cell fixation using trichloroacetic acid (TCA) and resolved by SDS-polyacrylamide gel electrophoresis before transfer to nitrocellulose membranes.

### Fractioning and Immunoblotting

For chromatin-enriched fractions around 6 × 10^7^ exponentially growing cells were harvested by centrifugation and resuspended in 1 ml of Buffer 1 (containing 150 mM Tris pH 8.8, 10 mM dithiothreitol (DTT), and 0.1% sodium azide), and incubated at room temperature for 10 minutes. Cells were pelleted, washed with 1 ml of Buffer 2 (50 mM KH_2_PO_4_/K_2_HPO_4_ pH 7.4, 0.6 M Sorbitol, and 10 mM DTT), resuspended in 200μl of Buffer 2 supplemented with 40 μg Zymolyase-100T and incubated at 37 °C for 10 minutes with intermittent mixing. The resulting spheroplasts were washed with 1 ml of ice-cold Buffer 3 (50 mM HEPES pH 7.5, 100 mM KCl, 2.5 mM MgCl, and 0.4 M Sorbitol), followed by resuspension and a 5-minute incubation in 100 μl of EBX buffer (50 mM HEPES pH 7.5, 100 mM KCl, 2.5 mM MgCl, 0.25% Triton100, 1 mM phenylmethylsulfonyl fluoride (PMSF), Protease inhibitor tablets (EDTA-free, Roche), Leupeptin 1 μg/ml, Pepstatin 2.5 μg/ml, and RNAse 10 μg/ml), with occasional mixing. Aliquots of 30 μl of these disrupted cell suspensions were collected as whole cell extract samples (WCE). Remaining volume was layered onto 70 μl of cold EBX-S buffer (EBX buffer supplemented with 30% Sucrose) and subjected to centrifugation at 12000 rpm for 10 minutes at 4 °C. Aliquots of 30 μl of the resulting supernatant layer (Chromatin-free fraction) were also collected. After discarding supernatant, chromatin pellets were washed with 200 μl of EBX-S buffer, resuspended in 70μl of EBX buffer supplemented with 0.5 μl of Benzonase, and incubated on ice for 15 minutes (Chromatin fraction). SDS-PAGE loading buffer was added to each fraction.

The different protein extracts were separated by SDS-polyacrylamide gel electrophoresis and transferred to nitrocellulose membranes. Antibodies used for detection are listed in the Resources Table and were visualized using ECL reagents (Amersham Pharmacia Biotech) and film. The levels of proteins bound to chromatin were quantified using Quantity One 1-D Analysis Software (BioRad) and normalized with their corresponding Histone H2B values. All data in the bar graphs are presented as means SD in triplicate. A two-way analysis of variance (ANOVA) test was used to determine the statistical significance.

### Protein interaction analysis

Cells expressing tagged or untagged (control) proteins were fixed with 1% formaldehyde and harvested. Chromatin extracts were prepared in a Lysis Buffer containing 50 mM Hepes pH 7.5, 140 mM NaCl, 1 mM EDTA, 1% Tritón-X100, 0.1% Na-deoxycholate, and supplemented with Antiproteolytic Cocktail using glass beads. Extracts were cleared by centrifugation; soluble protein fractions were discarded, and chromatin pellets were sheared by sonication. Chromatin extracts were clarified and tagged proteins were enriched by immunoprecipitation with specific Tag antibodies previously bound to Protein G Dynabeads (5 hours at 4°C). Then, antibody-bound Protein G Dynabeads and controls were extensively washed with lysis buffer, and elution was carried out in SDS-PAGE loading buffer. Immunoprecipitates were resolved by SDS-PAGE gels, transferred to nitrocellulose membranes and analysed with specific-HRP conjugated antibodies.

### Two-dimensional DNA gels (2D-gel analysis)

DNA samples for neutral-neutral two-dimensional gel electrophoresis were prepared and analyzed as described previously^60,75^. DNA was cut with the NcoI restriction enzyme, transferred to Hybond-XL (GE Healthcare) nitrocellulose membrane, and hybridized to radiolabeled probes spanning the *ARS305 and ARS306* origins of DNA replication. For each origin of replication tested, the specific probe corresponds to the following coordinates (retrieved from SGD): *ARS305* (39073-40557, Chr III) and *ARS306* (73001-73958, Chr III). Images were acquired using a Molecular Imager FX (BioRad) and different replication-associated DNA molecules were quantified using Quantity One 4.6 software (BioRad).

## Acknowledgments

We are grateful to members of 08 research group at the IBMCC for helpful discussions. We would like to particularly thank Professor Anne Donaldson and Takashi Kubota PhD (University of Aberdeen) for *pol30* mutant and ChVLig1 strains. We are also grateful to the National BioResource Project, NBRP Japan for the *cdc9-7* strain. This work was supported by the Spanish Ministry of Science (grants PID2019-109616GB-100 to A.B. and M.P.S. and PID2020-116003GB-100 to R.B.) and Junta de Castilla y León (grant SA103P20 to A.B). J.Z. was supported by a Predoctoral Fellowship from the Junta de Castilla y León (JCyL). S.M. was supported by a University of Salamanca Postdoctoral Fellowship and a MSCA Postdoctoral Fellowship (grant n° 101106007). E.A. was supported by a JCyL Postdoctoral Fellowship. A.B. and M.P.S. Institution is supported by the “Programa de Apoyo a Planes Estratégicos de Investigaciόn de Excelencia” cofunded by the Junta de Castilla y Leόn and the European Regional Development Fund (CLC-2017-01).

## Author Contributions

Conceptualization, A.B., with substantial inputs from M.P.S., J.Z., S.M. and RB; Investigation, J.Z., S.M., E.A., M.A. and M.P.S.; Supervision, A.B. and M.P.S.; Formal Analysis, A.B., M.P.S., S.M. and J.Z.; Writing & Editing, A.B. and M.P.S. with the help of R.B. and S.M.; Funding Acquisition, A.B., M.P.S. and R.B.

## Data availability

The datasets supporting the current study are available from the corresponding authors on request.

