## Supplemental Info (Table and 5 Figures) for "Timely lagging strand maturation relies on Ubp10-mediated PCNA dissociation from replicating chromatin"

The supplementary information is a single file that includes:

- A Resources Table containing yeast strains and reagents.
- Five Supplementary Figures (1-5) related to main Figures.

### Resources Table

| Figure | Strain | Genotype | Source |
|---|---|---|---|
| | | **All cells:** *W303 Mata ade2-1 can1- 100 his3-11,15 leu2-3,112 trp1-1 ura3-1 RAD5* | |
| **Figure 1** | 71.31 | *Mat a bar1::LEU2 Ubp10:GFP:kanMX* | Lab stock |
| **Figure 2A-C** | 55.34 | *Mat a bar1::LEU2* | Lab stock |
| | 73.64 | *Mat a bar1::LEU2 ubp10::hphMX* | Lab stock |
| | 84.63 | *Mat a bar1::LEU2 rad27::kanMX* | This study |
| | 84.70 | *Mat a bar1::LEU2 ubp10::hphMX rad27::kanMX* | This study |
| **Figure 2D** | 86.31 | *Mat a bar1::LEU2 Rad27:3Flag:HIS3* | This study |
| | 86.33 | *Mat a bar1::LEU2 Rad27:3Flag:HIS3 ubp10::hphMX* | This study |
| **Figure 3A** | 55.34 | As above | |
| | 87.53 | *Mat a bar1::HIS3 cdc9-7* | This study |
| | 73.64 | *As above* | |
| | 89.55 | *Mat a bar1::HIS3 cdc9-7 ubp10::natMX* | This study |
| **Figure 3B** | 55.34 | As above | |
| | 87.53 | As above | |
| | 89.55 | As above | |
| **Figure 3C-D** | 87.53 | As above | |
| | 89.55 | As above | |
| **Figure 4A-B** | 89.22 | *Mat a bar1::LEU2 Cdc9:3HA:kanMX* | This study |
| | 89.44 | *Mat a bar1::LEU2 Cdc9:3HA:kanMX Ubp10::natMX* | This study |
| **Figure 5A** | 87.53 | As above | |
| | 89.55 | As above | |
| | 87.72 | *Mat a bar1::hisG cdc9-7 pol30::LEU2:Pol30*$_{R14E}$ | This study |
| | 88.08 | *Mat a bar1::hisG cdc9-7 pol30::LEU2:Pol30*$_{R14E}$ *ubp10::kanMX* | This study |
| | 87.74 | *Mat a bar1::hisG cdc9-7 pol30::LEU2:Pol30*$_{D150E}$ | This study |
| | 88.34 | *Mat a bar1::hisG cdc9-7 pol30::LEU2:Pol30*$_{D150E}$ *ubp10::kanMX* | This study |
| **Figure 5B** | 87.53 | As above | |
| | 89.55 | As above | |
| | 88.08 | As above | |
| | 88.34 | As above | |
| **Figure 5C-D** | 87.57 | *Mat a bar1::hisG Ura3:pGal-OsTIR1 codon plus Cdc9-3xmini-AID:hphMX* | Donaldson Lab |
| | 87.60 | *Mat a bar1::hisG Ura3:pGal-OsTIR1 codon plus Cdc9-3xmini-AID:hphMX ubp10::kanMX* | This study |
| | 87.58 | *Mat a bar1::hisG Ura3:pGal-OsTIR1 codon plus Cdc9-3xmini-AID:hphMX pol30::LEU2:Pol30*$_{R14E}$ | Donaldson Lab |
| | 87.64 | *Mat a bar1::hisG Ura3:pGal-OsTIR1 codon plus Cdc9-3xmini-AID:hphMX pol30::LEU2:Pol30*$_{R14E}$ *ubp10::kanMX* | This study |
| | 87.59 | *Mat a bar1::hisG Ura3:pGal-OsTIR1 codon plus Cdc9-3xmini-AID:hphMX pol30::LEU2:Pol30*$_{D150E}$ | Donaldson Lab |
| | 87.66 | *Mat a bar1::hisG Ura3:pGal-OsTIR1 codon plus Cdc9-3xmini-AID:hphMX pol30::LEU2:Pol30*$_{D150E}$ *ubp10::kanMX* | This study |
| **Figure 6A-B** | 87.57 | As above | |
| | 87.60 | As above | |
| | 87.66 | As above | |
| **Figure 6C** | 87.53 | As above | |
| | 89.55 | As above | |
| | 88.34 | As above | |

| | | | |
|---|---|---|---|
| | 87.74 | As above | |
| **Figure 7A** | 55.34 | As above | |
| | 77.38 | *Mat a bar1::LEU2 elg1::kanMX* | Lab stock |
| | 79.65 | *Mat a bar1::LEU2 elg1::kanMX ubp10::natMX* | Lab stock |
| | 73.64 | As above | |
| **Figure 7B** | 73.64 | As above | |
| | 77.38 | As above | |
| | 79.65 | As above | |
| **Figure7C** | 55.34 | As above | |
| | 77.38 | As above | |
| | 79.65 | As above | |
| | 73.64 | As above | |
| **Figure S1A** | 81.09 | *Mat a Pol3:9PK:TRP1 ubp10:13Myc:hphMX ura3::URA3:GPD-TK7X* | Lab stock |
| **Figure S1B** | 85.16 | *Mat a bar1::LEU2 Ubp10:9PK:kanMX Cdc9:3Flag:HIS3* | This study |
| **Figure S1C** | 71.31 | As above | |
| **Figure S2** | 55.34 | As above | |
| **Figure S3A-B** | 55.34 | As above | |
| | 87.53 | As above | |
| | 89.55 | As above | |
| **Figure S4A** | 89.57 | *Mat a URA::pRS306-pGal1-10:OsTIR1 (pRS423-PGal1-10:HIS3)* | Kubota Lab |
| | 89.58 | *Mat a URA::pRS306-pGal1-10:OsTIR1 (pRS423-PGal1-10-ChVLig-3HA:HIS3)* | Kubota Lab |
| | 89.62 | *Mat a URA::pRS306-pGal1-10:OsTIR1 (pRS423-PGal1-10:HIS3) ubp10::kanMX* | This study |
| | 89.64 | *Mat a URA::pRS306-pGal1-10:OsTIR1 (pRS423-PGal1-10:HIS3) ubp10::kanMX* | This study |
| **Figure S4B-C** | 55.34 | As above | |
| | 73.64 | As above | |
| | 89.80 | *Mat a bar1::LEU2 ubp10::hphMX PGal-Cdc9-3HA:TRP1* | This study |
| **Figure S5A-B** | 55.34 | As above | |
| | 73.64 | As above | |
| | 77.38 | As above | |
| | 79.65 | As above | |
| **Figure S5C** | 88.70 | *Mat a bar1::LEU2 Elg1:3HA:URA3* | This study |
| | 89.40 | *Mat a bar1::LEU2 Elg1:3HA:URA3 ubp10::kanMX* | This study |

| Reagent or Resource | Source | Identifier |
|---|---|---|
| **Antibodies** | | |
| Anti-PCNA rabbit polyclonal (described) | Lab Stock | Generated at Home |
| Anti-H2B | Active Motif | Cat#39237; RRID:AB_2631110 |
| Anti-HA-HRP | Miltenyi Biotec | Cat#130-091-972; RRID:AB_ 871936 |
| Anti-PGK1 | Molecular Probes | Cat#A-6457; RRID:AB_221541 |
| Anti-Myc Tag (9E10) | Sigma-Aldrich | Cat#M5546; RRID:AB_260581 |
| Anti-Myc-HRP | Miltenyi Biotec | Cat#130-092-113; RRID:AB_871937 |
| Anti-PK V5 (E10/V4RR) | Thermo Fisher | Cat#MA5-15253; RRID:AB_2537639 |
| Anti-Flag M2 peroxidase (HRP) | Sigma-Aldrich | Cat#F1804; RRID:AB_259529 |
| Anti-rabbit IgG-Peroxidase | Amserham | Cat#NA934V; RRID:AB_772191 |
| Anti-mouse IgG-Peroxidase | Amserham | Cat#NA931V; RRID:AB_2721110 |
| **Chemicals Peptides and Recombinant Proteins** | | |
| Formaldehyde | Sigma-Aldrich | Cat#F8775 |
| GFP-Trap Agarose | Chromotek | Cat#GTA RRID:AB_2631357 |
| RNAse A | Sigma-Aldrich | Cat#10109169001 |
| Proteinase K | Sigma-Aldrich | Cat#3115852001 |
| Complete Protease Inhibitor-EDTA free | Sigma-Aldrich | Cat#11873580001 |
| Hydroxyurea | Ibian Technologies | Cat#HDU0250 |
| Alpha-factor Mating Pheromone | GenScript | Cat#RP01002 |
| Protran 0.45μM Nitrocelulose | Cytiva | Cat#GE10600003 |
| Spermine | Sigma-Aldrich | Cat#S1141 |
| Spermidine | Sigma-Aldrich | Cat#S2501 |
| MicroSpin G50 Columns | Cytiva | Cat#27533002 |
| Amersham Hybond-XL | GE Healthcare | Cat#RPN203S |
| DNA Polymerase I, Klenow Fragment | TakaraBio | Cat#2140A |
| Random Hexamers primer | Thermo Fisher | Cat#SO142 |
| T4 DNA Ligase | New England BioLabs | Cat#M0202S |
| SITOX Green | Invitrogen | Cat#S7020 |
| DynaBeads Protein-G | Invitrogen | Cat#10004D |
| **Deposited data** | | |
| Proteomic Ubp10 data | This study | ProteomeXchange: PXD048249 |
| **Oligonucleotides** | | |
| 5´ GGAGTTGGCCACGCTCTGGC 3´*ARS305* F | This study | N/A |
| 5´ CGCAACTACCCTAGAGCCTCTCCGCC 3´ *ARS305* R | This study | N/A |
| 5´ GGAAAAAAAGAAGACAAAG 3´ *ARS306* F | This study | N/A |
| 5´ CTTTAACTGCTCGTCGTAC 3´*ARS306* R | This study | N/A |

### SUPPLEMENTARY FIGURE LEGENDS

Supp Figure 1
Zamarreño et al., 2024

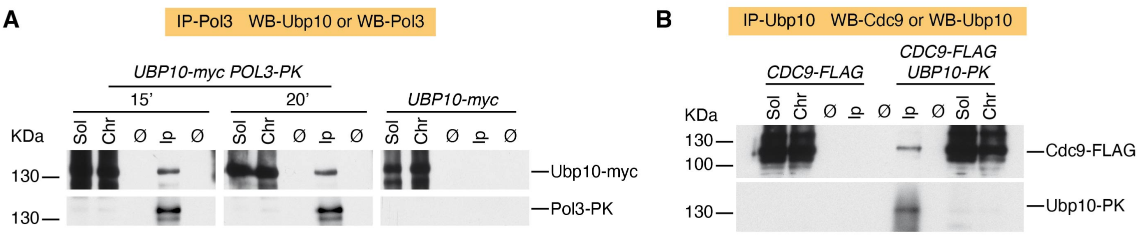

**C**

**S-phase Ubp10 Proteome (Ubp10-GFP-trap)**

| Selected identified Interactors | Gene names | Score | Unique peptides |
|---|---|---|---|
| RNA polymerase I subunit | *RPA190* | 323 | 37 |
| RNA polymerase I subunit | *RPA34* | 30 | 6 |
| RNA polymerase I subunit | *RPA43* | 70 | 9 |
| RNA polymerase I subunit | *RPA49* | 136 | 18 |
| RNA polymerase I subunit | *RPA135* | 307 | 34 |
| FACT complex subunit | *SPT16* | 183 | 34 |
| FACT complex subunit | *POB3* | 110 | 13 |
| Ubiquitin hydrolase 6 | *UBP6* | 63 | 12 |
| Ubiquitin hydrolase 14 | *UBP14* | 24 | 5 |
| Ubiquitin hydrolase 10 | *UBP10* | 323 | 35 |

**Supplementary Figure 1 (related to Figure 1). PCNA-DUB Ubp10 interacts in early S phase with the catalytic subunit of Pol∂ (Pol3) and DNA ligase I (Cdc9). A.** Coimmunoprecipitation assay showing physical interaction between Ubp10-myc and PK tagged Pol3. Pol3-PK was immunoprecipitated from formaldehyde-crosslinked protein extracts from untreated S phase cells, blots were incubated with $\alpha$-myc (to detect Ubp10) or $\alpha$ -PK (to detect Pol3). The immunoblots shown are those from alpha-factor synchronized cells and released into fresh media. Samples were taken at the indicated time points and tested by immunoblot. A *POL3* untagged strain was used as negative control. Soluble (Sol), chromatin (Chr) fraction control extracts and immunoprecipitates (Ip) are indicated. **B.** ChIP-CoIP analysis of Ubp10 interaction with the DNA ligase Cdc9. Coimmunoprecipitation analysis showing physical interaction between Ubp10-PK and Cdc9-FLAG in S phase. Cultures of *ubp10*-PK *cdc9*-FLAG cells, pre-synchronized in G1 with $\alpha$-factor, were released into S phase in fresh YPAD medium. Samples were taken 15 minutes after the G1 release and analyzed by Co-IPs and immunoblot. Soluble (Sol) and chromatin (Chr) fraction extracts and mock Ip controls are indicated as well as Ips. A *UBP10* untagged strain was used as negative control. Experiments in A and B were repeated 2 and 3 times respectively. **C**. RNA polymerase I complex, and FACT complex subunits were identified in the Ubp10-GFP-Trap IPs experimental approach subjected to LC-MS-MS analysis as described in figure 1.

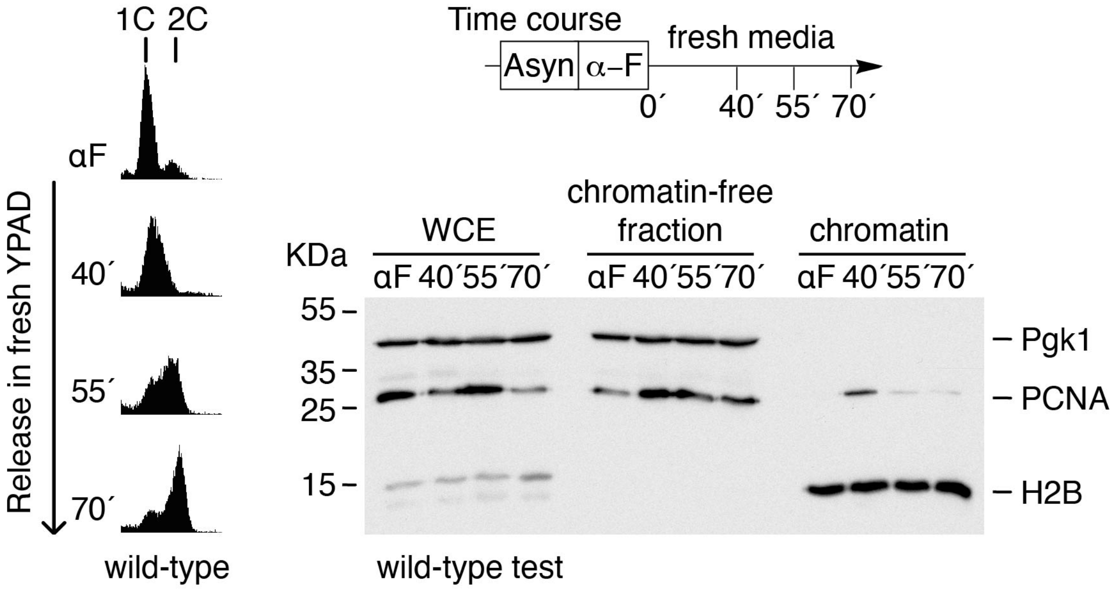

**Supplementary Figure 2 (related to Figures 2, 3, 4, 6 and 7). S phase chromatin association of PCNA in wild-type cells.** A log culture on unperturbed wild-type cells was synchronized in G1 with α-factor pheromone and released in fresh media to test S phase chromatin association of PCNA at indicated intervals. Whole cell extracts (WCE), chromatin-free fractions and chromatin-enriched fractions of each sample were prepared and electrophoresed in SDS-PAGE gels. Blots were incubated with α-Pgk1, α-PCNA or α-H2B antibodies. Pgk1 was used as a cytoplasmic control protein and histone H2B as a chromatin-associated control protein. DNA content analysis by FACS is shown on the left.

Supp Figure 3 Zamarreño et al., 2024

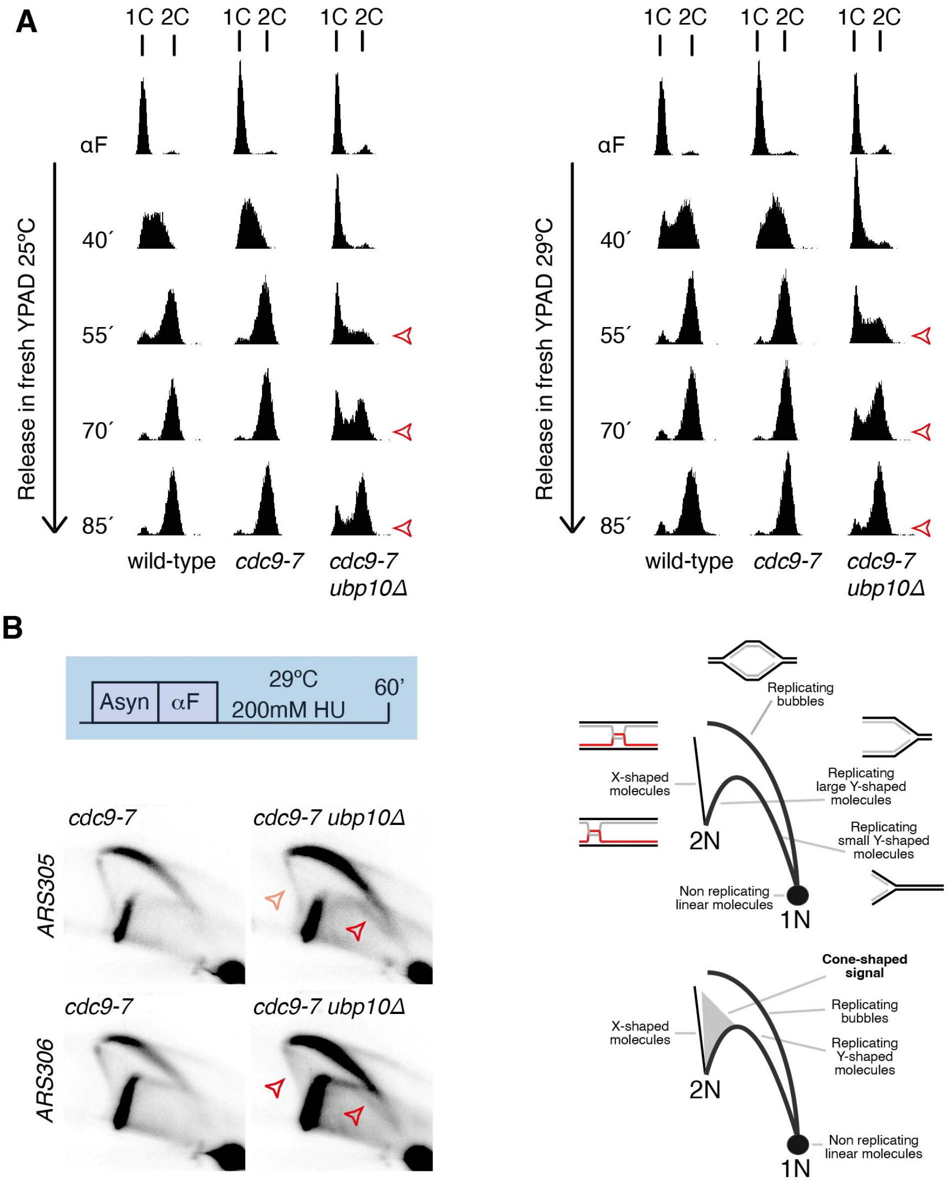

**Supplementary Figure 3 (related to Figure 3). S phase progression analysis of *cdc9-7 ubp10*Δ mutant cells. A.** DNA replication progression defects in *cdc9-7 ubp10*Δ cells both at 25ºC and 29ºC. DNA content analysis of wild-type, *cdc9-7,* and *cdc9-7 ubp10*Δ strains. Cells of the indicated strains were synchronized with α-factor and released in fresh yeast complex media (YPAD) either at 25ºC or 29ºC. **B**, Deletion of *UBP10* in *cdc9-7* leads to the accumulation of unusual replication intermediates. 2D-gel analysis of cells synchronized in early S phase with the ribonucleotide reductase inhibitor HU at 29ºC. Indicated strains were grown till exponential phase at 25ºC, synchronized in G1 with α-factor and released in fresh media with 200 mM HU at 29ºC. Under these conditions, *cdc9-7 ubp10*Δ double mutant cells accumulate abnormal DNA replication structures (cone-shaped signals and small Ys as indicated by red open arrows). A drawing of the normal replication intermediates and of the abnormal intermediates related to DNA replication fork collapse is shown.

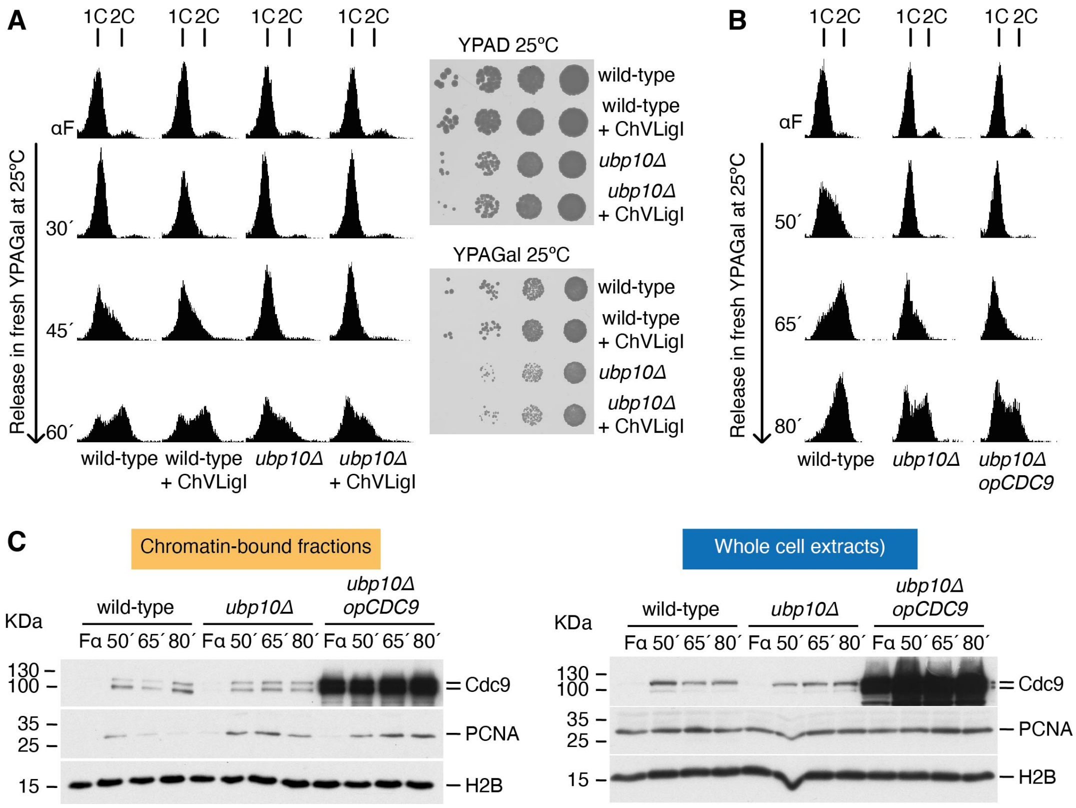

**Supplementary Figure 4 (related to Figure 4). Chromatin-bound Cdc9 fails to rescue *ubp10*Δ slow S phase phenotype**. **A.** Overexpression of *Chlorella* virus DNA ligase does not rescue S phase or growth defects in *ubp10*Δ mutant cells. DNA content analysis of wild-type and *ubp10*Δ strains harboring a plasmid containing the *Chlorella* virus DNA ligase (*GAL1,10:* ChVLigI) and controls (as indicated). Cells of the different strains were cultured al 25ºC in selective media (-HIS+Galactose) overnight to mid log phase, pelleted and reinoculated in fresh YPAGalactose. Further incubated four hours at 25ºC, synchronized with α -factor and released in fresh YPAGalactose. Samples were taken at indicated intervals and processed for FACS analysis of DNA content. (On the right) 10-fold dilution analysis of the indicated strains in Glucose or Galactose to repress or induce the expression of *GAL1-10* controlled ChVLigI. B. DNA content analysis by FACS of strains used (and described) in C. **C.** Chromatin association of Cdc9 in wild-type, *ubp10*Δ and *GAL1-10:CDC9-HA ubp10*Δ cells (all *CDC9-HA* tagged). Exponentially growing cultures of the three indicated strains grown to exponentially phase in YPA-Raffinose media were synchronized with α-factor and released in YPA-Galactose fresh media to test S phase chromatin association of Cdc9 under overexpressing conditions. Samples were taken at indicated intervals (α-factor, 50, 65 and 80 minutes); chromatin-enriched fractions were prepared and electrophoresed in SDS-PAGE gels. Blot was cut (according to MW) and incubated with α-Ha (to detect Cdc9-Ha), α -PCNA or α-H2B antibodies. Note that Cdc9-Ha remains associated with chromatin all throughout the time course experiment in *GAL1-10:CDC9-HA ubp10*Δ cells (overexpressing Cdc9). Aliquot samples as in A were processed with TCA to test Cdc9-Ha protein amounts in whole cell extracts (WCE).

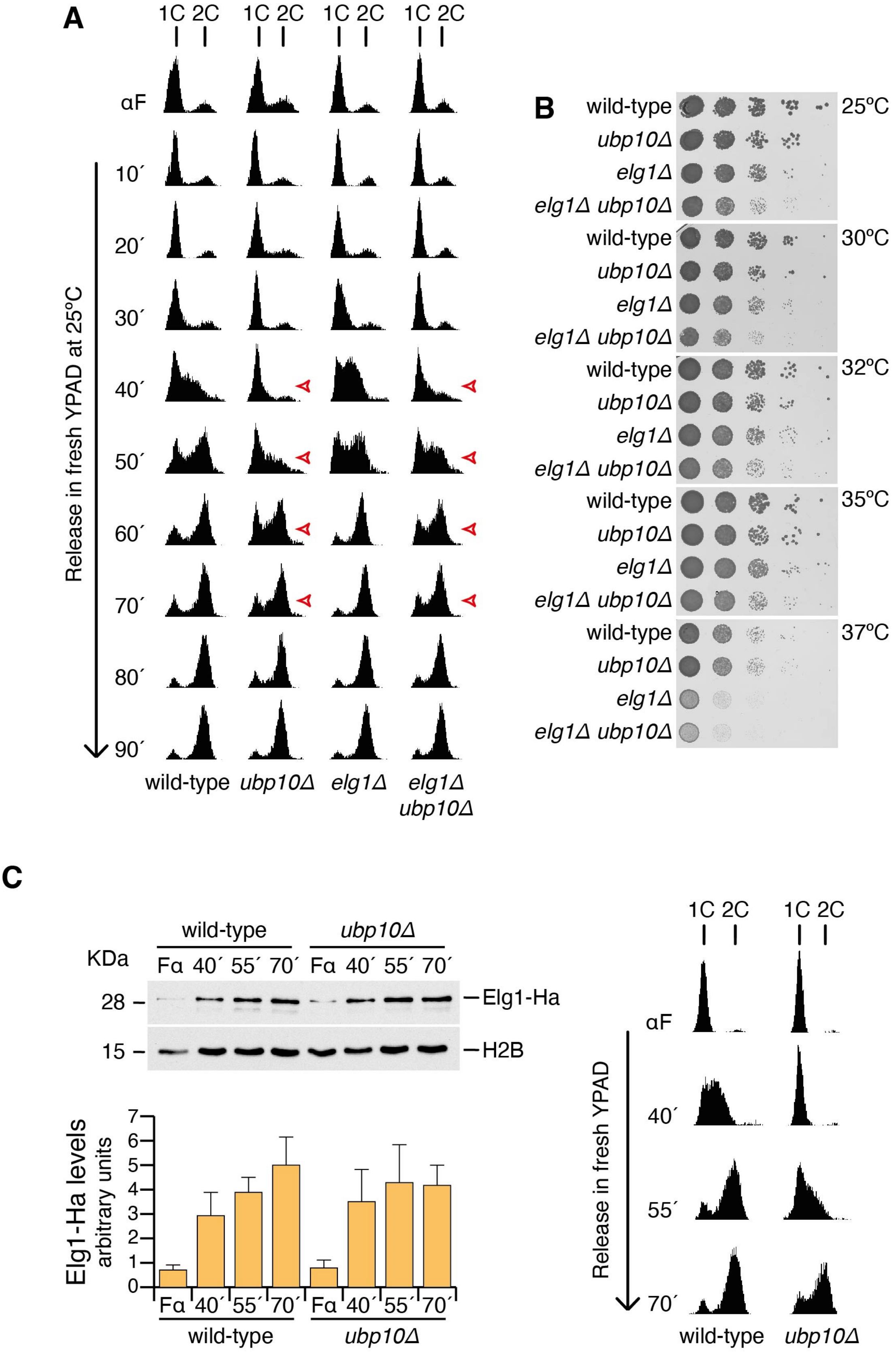
A
1C 2C
αF
10´
20´
30´
40´
50´
60´
70´
80´
90´
Release in fresh YPAD at 25ºC
wild-type
ubp10Δ
elg1Δ
elg1Δ ubp10Δ
B
wild-type
ubp10Δ
elg1Δ
elg1Δ ubp10Δ
25ºC
30ºC
32ºC
35ºC
37ºC
C
KDa
wild-type
ubp10Δ
Fα 40´ 55´ 70´ Fα 40´ 55´ 70´
28
15
Elg1-Ha
H2B
Elg1-Ha levels
arbitrary units
0
1
2
3
4
5
6
7
1C 2C
αF
40´
55´
70´
Release in fresh YPAD
wild-type
ubp10Δ

**Supplementary Figure 5 (related to Figure 7). *UBP10* is epistatic to *ELG1* in cell cycle progression but additive in growth rate. A.** *ubp10*Δ *elg1*Δ mutant cells progress slowly throughout S phase (similarly to *ubp10*Δ single mutants). DNA content analysis of wild type, *ubp10*Δ, *elg1*Δ and *elg1*Δ *ubp10*Δ strains. Cells of the indicated strains were synchronized with α-factor and released in fresh YPAD at 25ºC. The progression of the bulk genome replication was monitored at the indicated time points. Open red arrows indicate approximate S phase duration in *elg1*Δ and *elg1*Δ *ubp10*Δ strains. **B.** Poor growth rate in *ubp10*Δ *elg1*Δ mutant cells (as compared to single mutants or wild type cells), genetic interaction that points to semi-lethality. Ten-fold dilutions of wild type, *ubp10*Δ, *elg1*Δ and *elg1*Δ *ubp10*Δ strains incubated in rich media at indicated temperatures for 60 hours. Note the relative poor growth of the *elg1*Δ *ubp10*Δ strain. **C.** Elg1 binds to replicating chromatin in a Ubp10-independent manner. Chromatin association of Elg1 in wild-type and *ubp10*Δ in S phase. Exponentially growing cultures of (*ELG1*-Ha-tagged) wild-type and *ubp10*Δ cells were synchronized in G1 with α-factor and released in fresh media to test S phase chromatin association of Elg1. Samples were taken at indicated intervals; chromatin-enriched fractions were prepared and electrophoresed in SDS-PAGE gels. Blots were incubated with α-Ha (to detect Elg1-Ha) or α-H2B antibodies. A blot from a representative experiment is shown. Data in the graph represent the average of three biological replicates (and is expressed as means ±SD in triplicate) ($p = 0.5258$, two-way ANOVA test).
